# Oculopharyngeal muscular dystrophy (OPMD) associated alanine expansion impairs the function of the nuclear polyadenosine RNA binding protein PABPN1 as revealed by proximity labeling and comparative proteomics

**DOI:** 10.1101/2025.05.23.655706

**Authors:** Allison T. Mezzell, Alexandra M. Perez, Yu Zhang, Katherine E. Vest

**Affiliations:** Department of Molecular and Cellular Biosciences, University of Cincinnati

## Abstract

Oculopharyngeal muscular dystrophy (OPMD) is a late-onset disease caused by modest alanine expansion at the amino terminus of the nuclear polyadenosine RNA binding protein PABPN1. PABPN1 is expressed ubiquitously and is involved in multiple steps in RNA processing including optimal cleavage and polyadenylation, polyadenylation signal selection, and export of polyadenylated RNAs from the nucleus. Expanded PABPN1 forms aggregates in a subset of muscle nuclei, but PABPN1 levels are paradoxically low in muscle compared to other tissues. Despite several studies in model systems and patient tissues, it remains unclear whether alanine expansion directly impairs PABPN1 function. The molecular mechanisms leading to OPMD pathology are poorly understood. Here we used a proximity labeling approach to better understand the effect of alanine expansion on PABPN1 function in a cell culture model of skeletal muscle. To avoid the confounding factor of overexpression, PABPN1 constructs containing a carboxy-terminal TurboID tag were expressed in skeletal myotubes at near native levels using an inducible promoter. Although non-expanded PABPN1-TurboID was able to complement RNA export and myoblast differentiation defects caused by deficiency of endogenous PABPN1, alanine expanded PABPN1-TurboID was not. Comparative proteomics revealed increased interaction between expanded PABPN1 and RNA splicing and polyadenylation machinery and follow-up studies identified a dominant negative effect on RNA export in differentiated myotubes. These data indicate that alanine expansion can impair PABPN1 function regardless of the presence of wild type PABPN1 and support a model wherein both loss function and dominant negative effects of expanded PABPN1 contribute to OPMD pathology.

**Author summary:** Oculopharyngeal muscular dystrophy (OPMD) is a late onset muscle disease that, unlike other muscular dystrophies, causes progressive weakness primarily in muscles of the face and head. OPMD is most commonly inherited in a dominant fashion and is caused by small expansions in the gene encoding the RNA binding protein PABPN1. Despite the fact that PABPN1 is expressed ubiquitously and contributes to multiple steps in RNA processing, its expansion causes disease almost exclusively in skeletal muscle. The precise molecular events leading to OPMD disease onset have been difficult to characterize as it is challenging to separate functions expanded versus non-expanded PABPN1 in muscle without the confounding factor of overexpressing tagged fusion proteins. Here we fused expanded and non-expanded PABPN1 to the modified biotin ligase TurboID to label proximal proteins. PABPN1-TurboID fusions were expressed in a cell culture model of skeletal muscle at near-native levels. Using these constructs, we discovered that expanded PABPN1 is less functional than non-expanded PABPN1 and causes dominant negative effects on some PABPN1 functions. These findings support a model where both loss of function and dominant negative effects contribute to the molecular pathology of OPMD.

## Introduction

Oculopharyngeal muscular dystrophy (OPMD) is a late onset disease that causes progressive weakening of craniofacial and proximal limb muscles^1^. Although OPMD is rare globally (1:100,000), some populations are affected at rates up to 1:600^2^. The overwhelming majority of OPMD cases are caused by a dominant stable expansion of *GCN* codons in the gene encoding the <u>p</u>olyadenosine [poly(A)] RNA <u>b</u>inding <u>p</u>rotein <u>n</u>uclear 1 (PABPN1)^3^. Expansion of the PABPN1 protein occurs within an amino (N) terminal 10-residue alanine tract resulting in the addition of 1-8 alanine residues^1,4,5^. PABPN1 performs multiple functions in RNA processing including facilitating RNA poly(A) tail addition^6,7^, regulating polyadenylation signal (PAS) selection^8,9^, and promoting nuclear export of polyadenylated RNAs^10^. In the cytosol, PABPN1 remains bound to polyadenylated transcripts but the dominant species of poly(A) binding proteins is the cytoplasmic poly(A) binding protein PABPC1^1^. PABPN1 also participates in other steps of RNA processing including RNA decay^11^, terminal intron splicing^12^, and regulation of transcripts containing retained introns^13^. Although PABPN1 is expressed ubiquitously, the presence of expanded PABPN1 causes dysfunction in a small subset of skeletal muscles^2^. Expanded PABPN1 forms nuclear aggregates that can sequester PABPN1, other proteins, and RNA in a small proportion of myonuclei^14,15^. Paradoxically, PABPN1 levels in skeletal muscle are low relative to other tissues, suggesting that muscle is sensitized to PABPN1 perturbation^16,17,18^. Whether alanine expansion directly impairs PABPN1 function is not known and the mechanistic details underlying OPMD pathology are poorly understood.

The pathologic effects of expanded PABPN1 expression have been explored in multiple model systems. The most commonly used transgenic murine model of OPMD expresses PABPN1 expanded to 17 alanine residues (Ala17) under the control of a skeletal muscle actin promoter (A17.1 mouse)^19^. These mice express Ala17 PABPN1 at levels 10-30 times higher than endogenous PABPN1 in skeletal muscle and display severe muscle pathology correlated with the presence of PABPN1 aggregates in ∼25% of myonuclei^19,20^. Muscles in A17.1 mice also exhibit widespread shifts in PAS selection similar to the shift caused by *PABPN1* deficiency in U2OS osteosarcoma cells^8,9^. Our previous study of skeletal muscle in a knock-in mouse model of OPMD that expresses Ala17 PABPN1 at native levels (*Pabpn1^+/A17^*) showed mild shifts in PAS utilization and reduced poly(A) tail length similar to a hemizygous *Pabpn1* knockout mouse, suggesting that alanine expansion may indeed impair PABPN1 function^21^. However, in studies exploring genetic strategies to treat OPMD, expression of exogenous non-expanded PABPN1 alone was not sufficient to reverse all signs of muscle dysfunction unless it was coupled with shRNA knockdown of expanded PABPN1^22^. These studies suggest that loss of PABPN1 function alone is not sufficient to drive OPMD related pathology. Given that wild type and expanded PABPN1 are nearly identical in size, electrophoretic mobility, and recognition by antibodies, mechanistic studies of expanded PABPN1 biochemistry have been rarely performed without using overexpression of tagged exogenous PABPN1. Given that overexpression of wild type (Ala10) PABPN1 alone is sufficient to cause muscle phenotypes^1,23^, its overexpression is a confounding factor that must be tightly controlled. Studies of endogenous PABPN1 function in skeletal muscle, where levels are exceedingly low^16–18^, are rarer still.

PABPN1, like other RNA binding proteins, performs its functions in complex with other proteins. Thus, experimental approaches to identify the effect of alanine expansion on PABPN1 binding partners may provide insight into the functional effects of alanine expansion on PABPN1. Previously, we used tagged PABPN1 constructs to perform immunoprecipitation and comparative proteomics to examine PABPN1 and expanded PABPN1 binding proteins in murine limb muscle^24^. This study identified a preferential interaction between the frontotemporal dementia/amyotrophic lateral sclerosis (FTD/ALS) associated splicing protein TDP-43. Expression of expanded PABPN1 lead to intron retention in the transcript encoding sortillin-1 (SORT1) and resulted in loss of SORT1 function. Though this study was one of the first to demonstrate specific functional consequences of expanded PABPN1 expression, the comparative proteomics experiments did rely on potentially confounding PABPN1 overexpression. More recently, two studies using proximity labeling constructs (BirA*) demonstrated overlapping binding partners between PABPN1 and another poly(A) RNA binding protein, ZC3H14^13,25^. These studies revealed antagonistic functions of the two proteins in exosome-mediated decay versus nuclear export of transcripts containing retained introns. Similarly, in a study demonstrating the importance of PABPN1 for terminal intron splicing TurboID was recently used to detect widespread interactions between PABPN1 and components of the splicing machinery in HeLa cells^12^. However, these studies were performed in non-muscle cells and did not compare expanded and non-expanded PABPN1.

Here, we report the effect of alanine expansion on PABPN1 function using proximity labeling in differentiated C2C12 myotubes as a model of mature skeletal muscle. We generated stable C2C12 cell lines encoding non-expanded (Ala10) and expanded (Ala17) PABPN1 constructs tethered to an optimized biotin ligase (TurboID)^26^ and expressed under the control of an inducible promoter. *Pabpn1* knockdown experiments indicated that Ala10 but not Ala17 PABPN1-TurboID was able to fully complement endogenous PABPN1 deficiency. Proteomic analysis revealed that, when expressed at near-native levels, Ala10 and Ala17 PABPN1-TurboID constructs labeled similar cohorts of proximal proteins. However, label-free quantification showed that PABPN1 proximal proteins detected at higher levels in Ala17 PABPN1-TurboID myotube lysates were enriched in components related to RNA splicing, polyadenylation, and the ubiquitin-proteasome system. We found that the Ala17-PABPN1 protein was more insoluble and degraded faster, and it showed reduced interaction with components of the nuclear transcription export (TREX) complex. Fluorescent in situ hybridization (FISH) using oligo-deoxythymidine [oligo d(T)] nucleotide probes suggested that bulk polyadenylated RNA is retained in the nucleus in C2C12 myotubes expressing Ala17 PABPN1-TurboID and in primary myotubes generated from the *Pabpn1^+/A17^* mouse model of OPMD. Taken together, these data suggest that expanded PABPN1 undergoes increased turnover and reduced interaction with nuclear export machinery leading to reduced export of polyadenylated RNAs. These results support a model in which expression of expanded PABPN1 leads to a dominant negative effect including phenotypes that resemble loss of PABPN1 function.

## Results

### Generation of stable cell lines encoding inducible PABPN1-TurboID constructs

PABPN1 is a low abundance protein in skeletal muscle and PABPN1 levels decrease during myoblast differentiation to myotubes (Figure S1A-C)^16,17^. To avoid the confounding effects of PABPN1 overexpression^1,23^, we sought to generate an experimental system expressing near-native levels of PABPN1 proximity labeling constructs. Lentiviral plasmids encoding non-expanded (Ala10) or expanded (Ala17) PABPN1 fused to a carboxy (C) terminal TurboID proximity labeling protein^26^ along with DYKDDDDK (Flag) and Myc tags under the control of the doxycycline-inducible TRE promoter were transduced into C2C12 myoblasts (Figure 1A). To determine if stable cells could be used to express near native levels of PABPN1-TurboID, transduced myoblasts were differentiated to myotubes, titrated with doxycycline, and treated with biotin (Figure 1A). Considering that expanded PABPN1 is expressed from one of two alleles in autosomal dominant OPMD, we defined near-native expression of PABPN1-TurboID as approximately 50% of endogenous PABPN1. Immunoblots probed with an antibody to BirA (the parent protein of engineered TurboID) (Figure 1B) or an antibody to the Myc tag (Figure S1D) showed dose-dependent expression of PABPN1-TurboID constructs in doxycycline-treated myotubes that is optimal at ∼48 hours of doxycycline treatment. Probing immunoblots with an antibody to PABPN1 revealed that Ala17 PABPN1-TurboID and the corresponding Ala10 PABPN1-TurboID control can be expressed at ∼50% of endogenous PABPN1 levels depending on dose of doxycycline (Figure 1C, D). Multiple bands were detected in blots of lysates from PABPN1-TurboID expressing myotubes when probed with streptavidin-conjugated horseradish peroxidase (HRP) in a doxycycline dose-dependent manner (Figure 1B), indicating that the TurboID fusions can adequately label multiple proteins in myotubes. A similar result was detected in blots of eluates from streptavidin bead capture (Figure 1E and S1E). Importantly, expression of Ala17 PABPN1-TurboID was confirmed in immunoblots of streptavidin elutions from induced myotubes using an antibody targeting the alanine expansion (Figure 1E and S1E)^27^. Taken together, these data indicate that PABPN1-TurboID constructs can be expressed at near-native levels in C2C12 myotubes.

**Figure 1:**
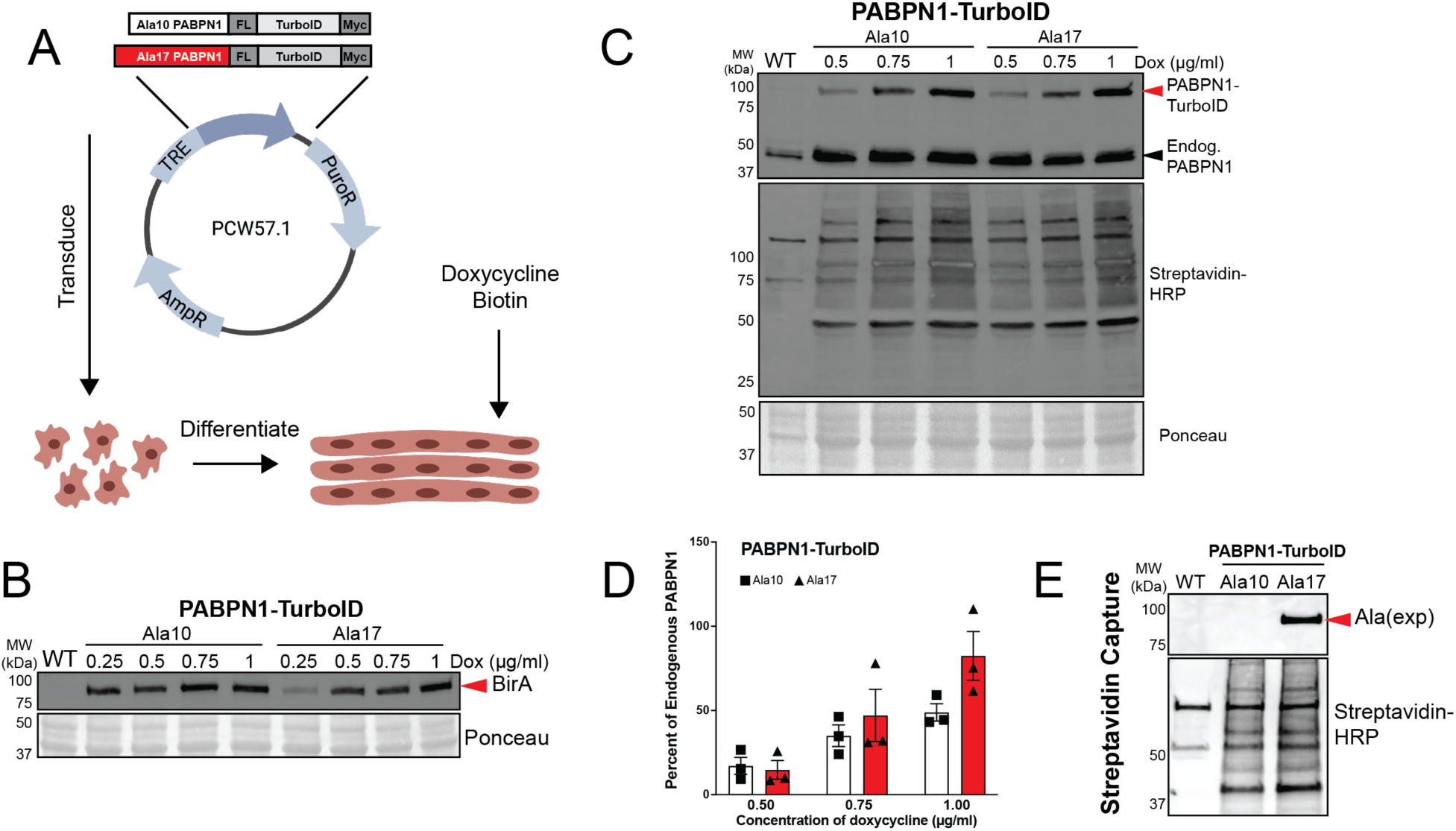
Ala10 and Ala17 PABPN1-TurboID constructs can be induced to express at near-native levels. **A)** Schematic outlining the experimental approach to express Ala10 and Ala17 PABPN1-TurboID in differentiated C2C12 myotubes. Genes encoding PABPN1-TurboID constructs containing Flag tag (FL) and Myc tag were cloned into the pCW57.1 lentiviral vector under the control of a tetracycline responsive element (TRE). C2C12 myoblasts were transduced and allowed to differentiate to myotubes. Lysates from doxycycline-treated myotubes were assayed by immunoblots shown and quantified in *B-D*. In all cases, Ponceau staining was used as a loading control. **B)** PABPN1-TurboID constructs detected as a band just below 100 kDa using an antibody to BirA, the parent protein of engineered TurboID. **C)** PABPN1-TurboID constructs (top band) as detected by antibody to PABPN1 (top blot) allowing for comparison to endogenous PABPN1 (bottom band) and detection of dose-dependent biotinylation using streptavidin conjugated to horseradish peroxidase (HRP, middle blot). **D)** Ratio of PABPN1-TurboID constructs to endogenous PABPN1 showing that PABPN1-TurboID can be expressed at near-native levels (∼50-100% of endogenous PABPN1). Shown are individual measurements and mean ± standard deviation for n = 3 experiments. **E)** Confirmation of expression of Ala17 PABPN1-TurboID as detected by an antibody to alanine expansion and of proximal protein biotinylation as detected by streptavidin-HRP in streptavidin capture eluates from doxycycline-treated myotubes. Shown is a blot from one representative elution with additional replicates shown in Supplementary Figure 1E. Wild type (WT) C2C12 myotubes were used as a control for endogenous biotin-binding proteins.

### Functional characterization of PABPN1-TurboID constructs

To verify that fusion of the TurboID protein to the PABPN1 C-terminus does not cause adverse effects, we sought to confirm the localization and functionality of PABPN1-TurboID constructs. Although many canonical PABPN1 functions occur in the nucleus, the PABPN1 protein is found in the nucleus and the cytoplasm^28^. Subcellular fractionation and immunoblot indicated that PABPN1-TurboID can be detected in both nuclear and cytoplasmic compartments in myotubes (Figure 2A). PABPN1-TurboID constructs were more prominent in the cytosol than endogenous PABPN1, and quantification showed a similar distribution ratio between Ala10 and Ala17 PABPN1-TurboID (Figure 2B). Notably, expression of PABPN1-TurboID did not have any effect on the distribution of endogenous PABPN1 (Figure 2C). Immunofluorescence staining of induced myotubes using an antibody to PABPN1 revealed that total PABPN1 (representing endogenous and TurboID constructs) was distributed normally with most signal in the nucleus and some signal in the cytosol (Figure 2D and S2A). Fluorescence detection of biotinylated proteins using AlexaFluor (AF)-488 conjugated streptavidin indicated a marked increase of biotinylated proteins primarily in the nuclei of myotubes expressing PABPN1-TurboID, while endogenous biotinylation in wild type control cells was detected primarily in the cytosol (Figure 2E and S2B). These data show that both Ala10 and Al7 PABPN1-TurboID constructs largely localize normally, though some additional PABPN1-TurboID can be detected in the cytosol.

**Figure 2:**
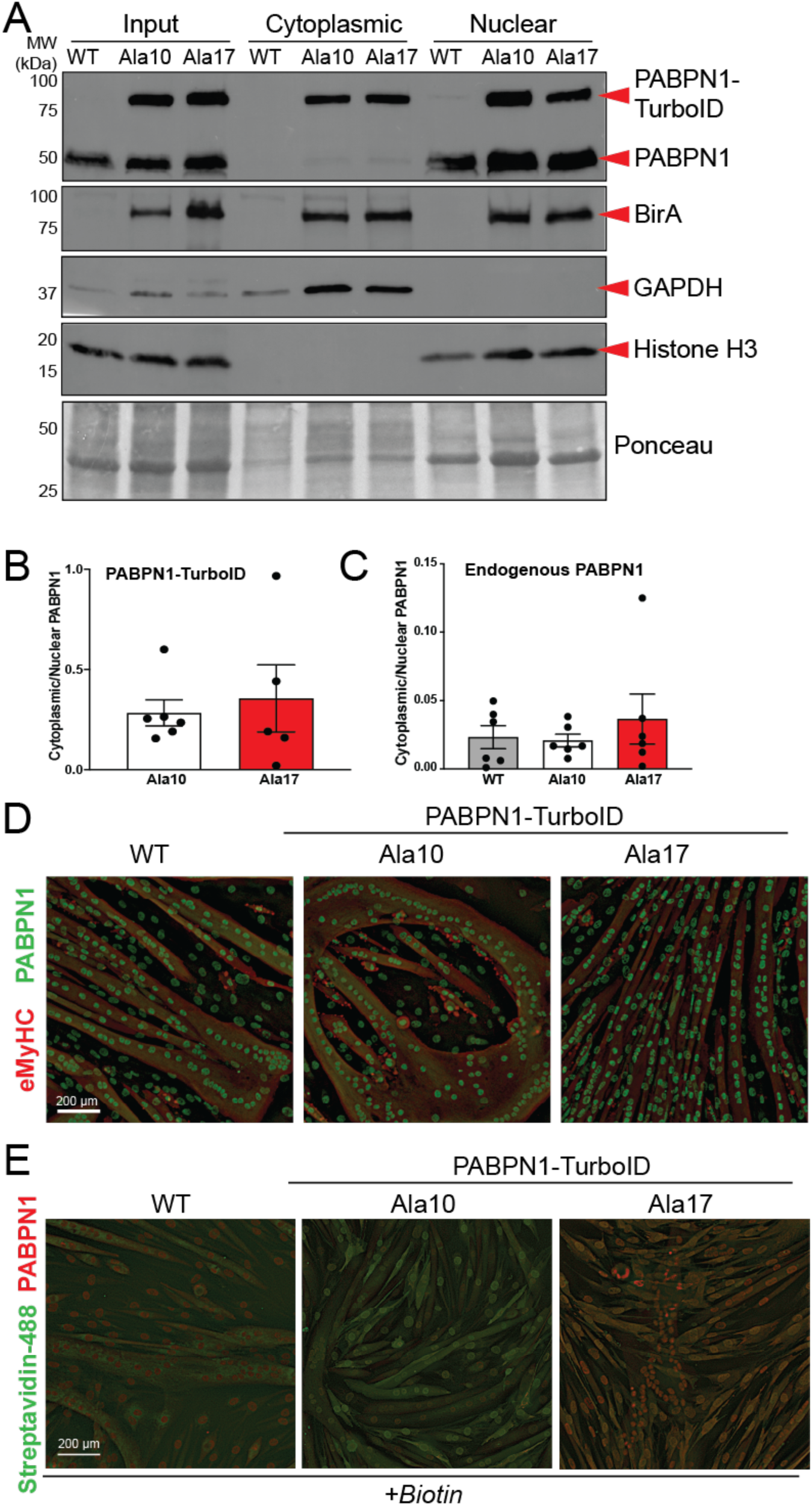
Ala10 and Ala17 PABPN1-TurboID constructs localize to the nucleus and cytoplasm in myotubes. **A)** Sub-cellular fractionation of myotube lysates derived from WT or stable cells expressing Ala10 and Ala17 PABPN1-TurboID analyzed by immunoblot using antibodies to PABPN1 and BirA. GAPDH and Histone-H3 serve as controls for cytoplasmic and nuclear fractions, respectively. Ponceau staining was used as a loading control. **B)** Quantification of cytoplasmic/nuclear ratio of PABPN1-TurboID constructs as detected by antibody to PABPN1 shown in *A*. Shown is mean ± standard deviation for n = 5-6 experiments. Statistical calculations used unpaired t-test. **C)** Quantification of cytoplasmic/nuclear ratio of endogenous PABPN1 as detected by antibody to PABPN1 shown in *A*. Shown is mean ± standard deviation for n = 6 experiments. Statistical calculation used ordinary one-way ANOVA. **D)** Immunostaining with an antibody to PABPN1 (green) in myotubes from WT and stable cells expressing near-native levels of Ala10 and Ala17 PABPN1-TurboID. Immunostaining with an antibody to embryonic myosin heavy chain (red) was used to detect myotubes. **E)** Staining using Alexa Fluor 488-conjugated streptavidin (Streptavidin-488, green) to detect biotin and an antibody to PABPN1 (red) to detect total PABPN1. For *D* and *E*, shown are representative images with additional replicates shown in Supplementary Figure 2.

Loss of PABPN1 function leads to impaired myoblast differentiation and nuclear accumulation of polyadenylated RNAs^10^. To determine if PABPN1-TurboID constructs function normally, we performed rescue experiments in *Pabpn1* knockdown cells. PABPN1 deficiency was induced in C2C12 myoblasts using siRNA with or without re-expression of Ala10 or Ala17 PABPN1-TurboID. As expected, PABPN1 deficiency resulted in nuclear accumulation of polyadenylated RNAs in myoblasts as detected by fluorescence in situ hybridization (FISH) using oligo d(T) probes conjugated with AF-488 (Figure 3A and S3). Expression of Ala10 PABPN1-TurboID reversed the nuclear RNA accumulation phenotype in *Pabpn1* knockdown cells while expression of Ala17 PABPN1-TurboID did not (Figure 3A and S3). Interestingly, myoblast growth was significantly reduced upon expression of Ala17 PABPN1-TurboID in negative control (*siScr*) cells as detected by nuclei count (Figure 3B), which is consistent with the reduced proliferation detected in pharyngeal myoblasts from the *Pabpn1^+/A17^* knock-in mouse in vivo and in vitro^29^. After inducing differentiation, as expected, loss of PABPN1 (Figure 3C, F) impaired myotube formation (Figure 3D) as measured by fusion index (Figure 3E). Upon expression of Ala10 PABPN1-TurboID (Figure 3B), the differentiation phenotype in PABPN1 deficient cells was restored to wild type levels (Figure 3C, D), but expression of Ala17 PABPN1-TurboID (Figure 3F) only partially rescued myotube formation (Figure 3G, H). Interestingly, the fusion index was significantly reduced in non-treated or scrambled siRNA-treated C2C12 myotubes (no knockdown of endogenous *Pabpn1*) that were expressing Ala17 PABPN1-TurboID while expression of Ala10 PABPN1-TurboID did not cause the same dominant negative effect (Figure 3I). PABPN1 deficiency also impaired survival in myoblasts induced to differentiate as measured by reduced total nuclei, and this phenotype was rescued by expression of both Ala10 and Ala17 PABPN1-TurboID (Figure 3J). These data indicate that the Ala10 PABPN1-TurboID can functionally complement PABPN1 deficiency, but Ala17 PABPN1-TurboID can only partially replace endogenous PABPN1. These results suggest that C-terminal fusion of the TurboID protein does not negatively impact PABPN1, but alanine expansion impairs some PABPN1 functions.

**Figure 3:**
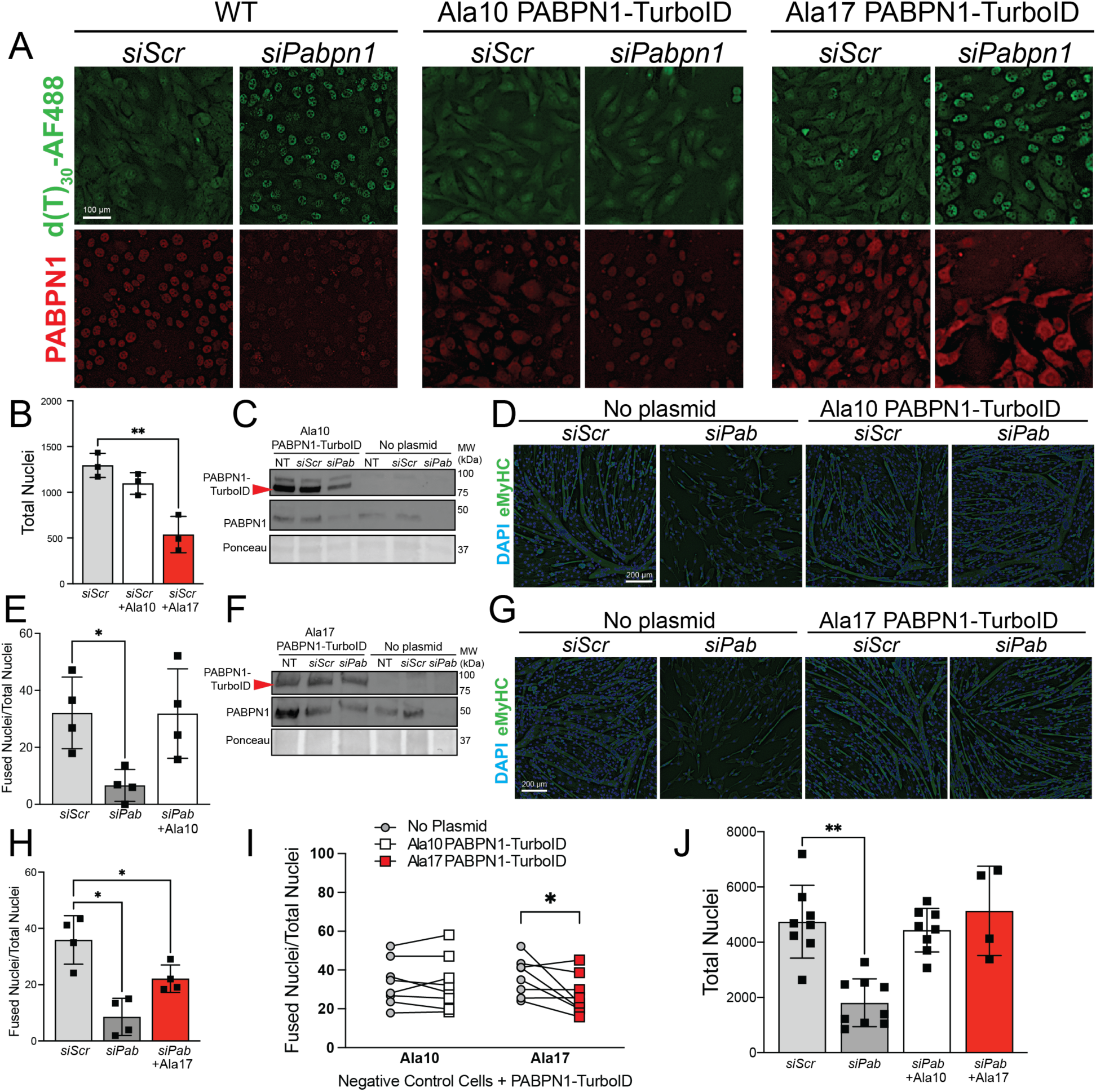
Ala10 PABPN1-TurboID functionally complements endogenous PABPN1 deficiency while Ala17 PABPN1-TurboID only partially rescues PABPN1 loss. **A)** Fluorescence in situ hybridization (FISH) in myoblasts with (*siPabpn1*) and without (*siScr*) *Pabpn1* knockdown using d(T)_30_-AF488 to detect polyadenylated RNA with immunostaining using an antibody to PABPN1 to show knockdown. FISH was performed in wild type (WT) cells or stable cells expressing near-native levels of Ala10 or Ala17 PABPN1-TurboID. Shown are representative images with additional replicates shown in Supplementary Figure 3. **B)** Quantification of total nuclei in control myoblasts (*siScr*) from *A* showing decreased cell count in *siScr* myoblasts expressing Ala17 PABPN1-TurboID. Shown is mean ± standard deviation for n = 3 experiments. Statistical significance was determined by one-way ANOVA. ** p < 0.01. **C)** Immunoblot probed with antibody to BirA (PABPN1-TurboID) or PABPN1 showing PABPN1 knockdown and replacement with Ala10 PABPN1-TurboID in myoblasts induced to differentiate in the presence of *Pabpn1* targeting siRNA (*siPab*) with or without co-transfection with plasmid encoding Ala10 PABPN1-TurboID. Ponceau staining was used as a loading control. **D)** Immunostaining using an antibody to embryonic myosin heavy chain (eMyHC) to visualize myotubes in cells with (*siPab*) and without (*siScr*) *Pabpn1* knockdown or expression of Ala10 PABPN1-TurboID. DAPI staining used to visualize nuclei. Shown are representative images. **E)** Fusion index calculated for *Pabpn1* knockdown myotubes (*siPab*) showing rescue of differentiation upon expression of Ala10 PABPN1-TurboID (+Ala10) in *Pabpn1* knockdown myotubes. Shown is mean ± standard deviation for n = 4 experiments. Statistical significance determined by one-way ANOVA. * p < 0.05. **F)** Immunoblot showing *Pabpn1* knockdown and replacement with Ala17 PABPN1-TurboID similar to experiment shown in *C*. **G)** Immunostaining using eMyHC antibody in *Pabpn1* knockdown and replacement with Ala17 PABPN1-TurboID similar to experiment shown in *D*. **H)** Fusion index similar to that shown in *E* showing only partial rescue of differentiation upon expression of Ala17 PABPN1-TurboID in *Pabpn1* knockdown myotubes. Shown is mean ± standard deviation for n = 4 experiments. Statistical significance determined by one-way ANOVA. * p < 0.05. **I)** Fusion index calculated for control myotubes (nontreated or *siScr*) expressing Ala10 or Ala17 PABPN1-TurboID showing dominant negative effect of Ala17 on fusion index. Shown are n = 8 replicates for each experiment. Statistical significance determined by two-way ANOVA. * p < 0.05. **J)** Total nuclei as a measurement of survival showing reduced nuclear count in *Pabpn1* knockdown myotubes (*siPab*) that is fully rescued upon expression of Ala10 or Ala17 PABPN1-TurboID. Shown is mean ± standard deviation for n = 4-8 experiments. Statistical significance determined by one-way ANOVA. ** p < 0.01.

### Identification of Ala10 and Ala17 PABPN1 proximal proteins using TurboID

To better understand how alanine expansion might impair PABPN1 function, we sought to identify proteins proximal to Ala10 and Ala17 PABPN1 using TurboID mediated biotin labeling^26^. Myotubes were induced to express near-native levels of Ala10 or Ala17 PABPN1-TurboID for 48 hours, treated with biotin for two hours, and lysates applied to streptavidin agarose beads. Elutions from streptavidin capture of biotinylated proteins were analyzed by SDS-PAGE. Silver staining and immunoblots indicated that multiple biotinylated proteins were captured from myotubes expressing PABPN1-TurboID (Figure S4 A and 1E). Importantly, immunoblot of streptavidin bead eluates revealed that both Ala10 and Ala17 PABPN1-TurboID labeled known PABPN1 binding partners PABPC1 and TDP-43 (Figure S4B). As expected, more TDP-43 was detected in streptavidin elutions from myotubes expressing Ala17 PABPN1-TurboID compared to those expressing Ala10 PABPN1-TurboID (Figure S4C)^24^. These data indicate that PABPN1-TurboID constructs biotinylate multiple proteins including known PABPN1 binding partners.

To identify additional binding partners of Ala10 and Ala17 PABPN1, proteomic analysis was performed on streptavidin capture of biotin-treated myotube lysates from wild type control cell lines and cell lines expressing near-native levels of Ala10 and Ala17 PABPN1-TurboID (Figure 4A). Comparative proteomic profiling was performed on streptavidin eluates from three independent experiments. A total of 595 proteins were identified with a false discovery rate (FDR) of 95% or higher (complete proteomics data can be found in Supplemental Table 2). To filter out potential contaminants, comparisons were performed on proteins that were detected in two of three independent experiments for Ala10 or Ala17 PABPN1-TurboID expressing myotubes but were not detected in wild type control samples. A total of 130 proteins were detected in Ala10 PABPN1-TurboID eluates and 142 proteins were detected in Ala17 PABPN1-TurboID eluates (Figure 4B). As expected, functional annotation of enriched gene ontologies (GO) and pathways using the STRING database^30^ revealed enrichment for terms associated with “RNA binding” and “metabolism of RNA” in both Ala10 and Ala17 PABPN1 proximal proteins (Figure 4C and S5A). Importantly, several known and putative PABPN1 binding proteins were identified in both Ala10 and Ala17 PABPN1-TurboID data sets including ZC3H14^31^, CPSF6^25,32^, NUDT21^25,32^, ZC3H11A^33^, RBM26 and RBM27^25^, and HNRNPA1^34^ (Supplemental Table 2).

**Figure 4:**
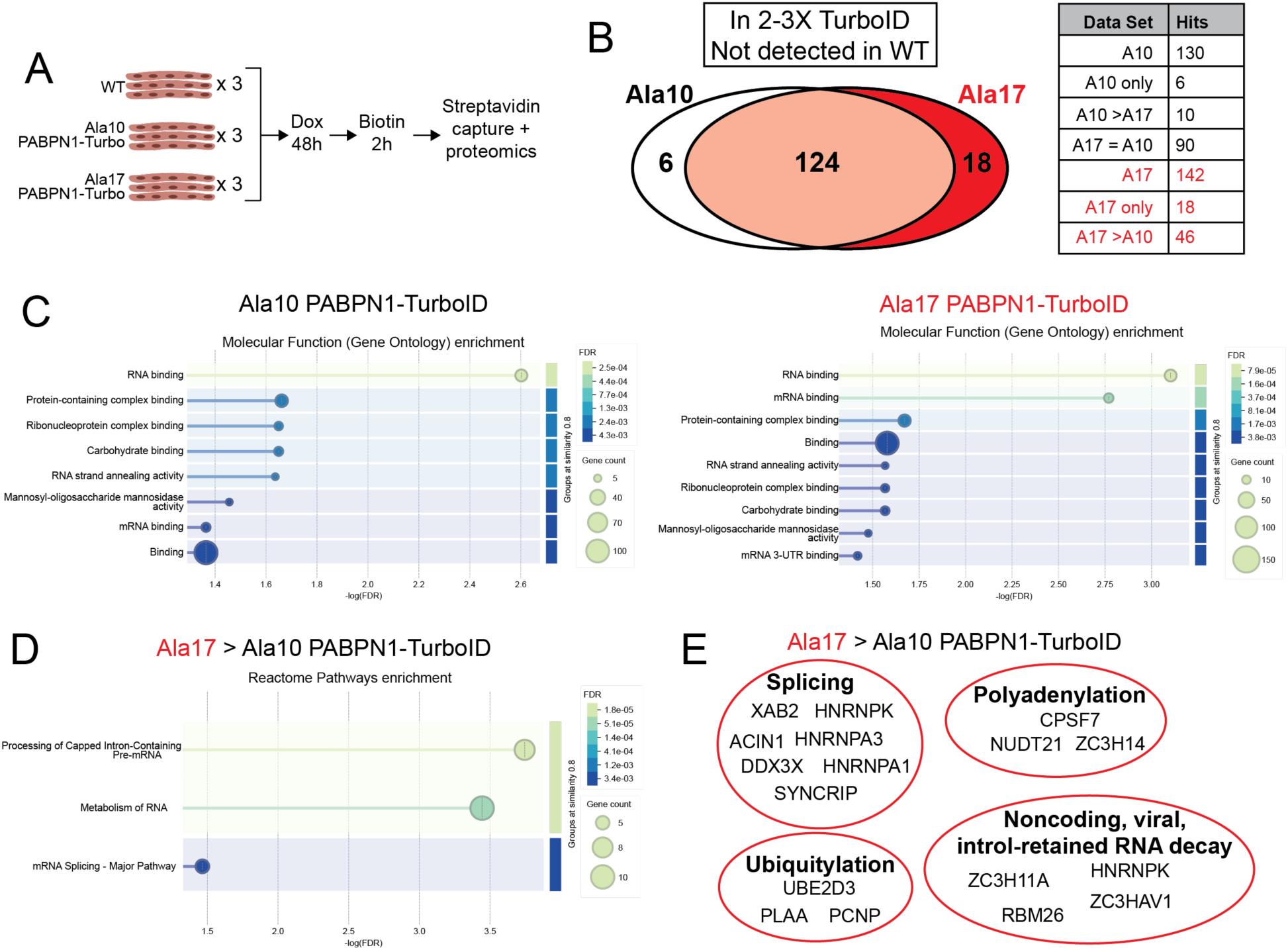
Proteomic analysis detects similar Ala10 and Ala17 PABPN1 proximal proteins and increased Ala17 PABPN1 interaction with splicing and polyadenylation machinery. **A)** Experimental design included three individual replicates each of wild type (WT) myotubes or myotubes expressing near-native levels of Ala10 or Ala17 PABPN1-TurboID. **B)** Proximal proteins detected in all three Ala10 or Ala17 PABPN1 replicates but not in WT samples revealed similar number putative interacting proteins for both constructs and 6 and 18 being unique to Ala10 and Ala17 PABPN1-TurboID, respectively. **C)** Gene ontology analysis showing similar molecular functions of Ala10 and Ala17 PABPN1-TurboID proximal proteins. **D)** Gene ontology analysis showing molecular function of Ala17 PABPN1 enriched proximal proteins were mostly related to RNA metabolism and RNA splicing. **E)** Proteins related to splicing, polyadenylation, RNA decay, and ubiquitylation enriched in Ala17 PABPN1 proximal proteins. Additional functional annotation shown in Supplementary Figure 5.

Other known PABPN1 interacting proteins like PABPC1 and TDP-43 were detected at high abundance in Ala10 and Ala17 PABPN1-TurboID samples but were also detected at low levels in wild type controls and were thus omitted from the functional annotation. Filtering proteomics data to include proteins detected in PABPN1-TurboID eluates at least two-fold higher than wild type controls yielded 324 and 346 hits for Ala10 and Ala17 PABPN1-TurboID, respectively (Figure S6A). Functional annotation of this more inclusive data set yielded similar GO and pathway enrichment as the more strictly filtered data set (Figure S6B). Several of these proteins were detected in published studies of PABPN1 binding partners^12,13,24,25,35^ (Supplemental Table 2). Thus, proteomic data indicate that Ala10 and Ala17 PABPN1-TurboID exist in close proximity to similar subsets of proteins.

To determine whether alanine expansion affects the magnitude of interaction of PABPN1 with its proximal proteins, we compared label-free quantification of proteomic data from streptavidin eluates of Ala10 versus Ala17 PABPN1-TurboID expressing myotubes. Proteins detected with fold change of one or more comparing Ala10 and Ala17 PABPN1-TurboID were considered differentially enriched. Of the proteins that were not detected in the wild type controls, ten were detected with higher abundance in the Ala10 PABPN1-TurboID samples and 46 were detected with higher abundance in Ala17 PABPN1-TurboID samples (Figure 4B). No significant enrichment of GO terms or pathways was detected in the proteins enriched in Ala10 PABPN1-TurboID data, but the list included multiple cytoplasmic proteins and one putative component of the CCR4-NOT deadenylation and decay complex (TNKS1BP1). Functional annotation of proteins enriched in Ala17 PABPN1-TurboID cells highlighted terms related to “RNA binding” and “processing of capped intron-containing pre-mRNA” (Figure 4D and S5B). Proteins enriched in the Ala17 PABPN1-TurboID interactome are involved in polyadenylation and cleavage (CPSF7, NUDT21, ZC3H14), RNA splicing (HNRNPA1, HNRNPA3, ACIN1, HNRNPK, XAB2, SYNCRIP) and decay of noncoding, intron-retaining, or viral RNAs (ZC3H11A, RBM26, ZC3HAV1, HNRNPK) (Figure 4E). A smaller group of proteins (UBE2D3, PLAA, PCNP) were related to the function of ubiquitylation and protein catabolism, which has been previously implicated in studies of OPMD pathology^36,37^. Similar pathways were identified in 77 Ala17 PABPN1-TurboID enriched proximal proteins when using the more inclusive data set of proteins detected at levels two-fold over wild type (Figure S6 C). Taken together, these data suggest that Ala17 PABPN1-TurboID exhibits increased interaction with proteins involved in RNA processing and proteasomal degradation, which may impact downstream PABPN1 functions.

### Effect of alanine expansion of PABPN1 function

Proteomics data suggest that Ala17 PABPN1-TurboID interacts more strongly with proteins related to protein catabolism including the E2 ubiquitin conjugating ligase UBE3D2 (Figure 4E, Supplemental Table 2). Previous studies using the *Pabpn1^+/A17^* knock-in mouse model of OPMD do not report any changes in steady-state levels of total PABPN1 in murine skeletal muscle but in that model wild type and alanine-expanded PABPN1 are biochemically indistinguishable^21,29^. Here, cycloheximide chase experiments were used to determine the effect of alanine expansion on soluble PABPN1 stability. Ala10 and Ala17 PABPN1-TurboID expressing myotubes were treated with cycloheximide to stop translation and lysates were harvested intermittently for 24 hours. Immunoblot analysis indicated that Ala10 PABPN1-TurboID and endogenous PABPN1 in wild type cells were largely stable up to ∼12 hours (Figure 5 A, B) while levels of Ala17 PABPN1-TurboID decreased relatively quickly (Figure 5C).

**Figure 5:**
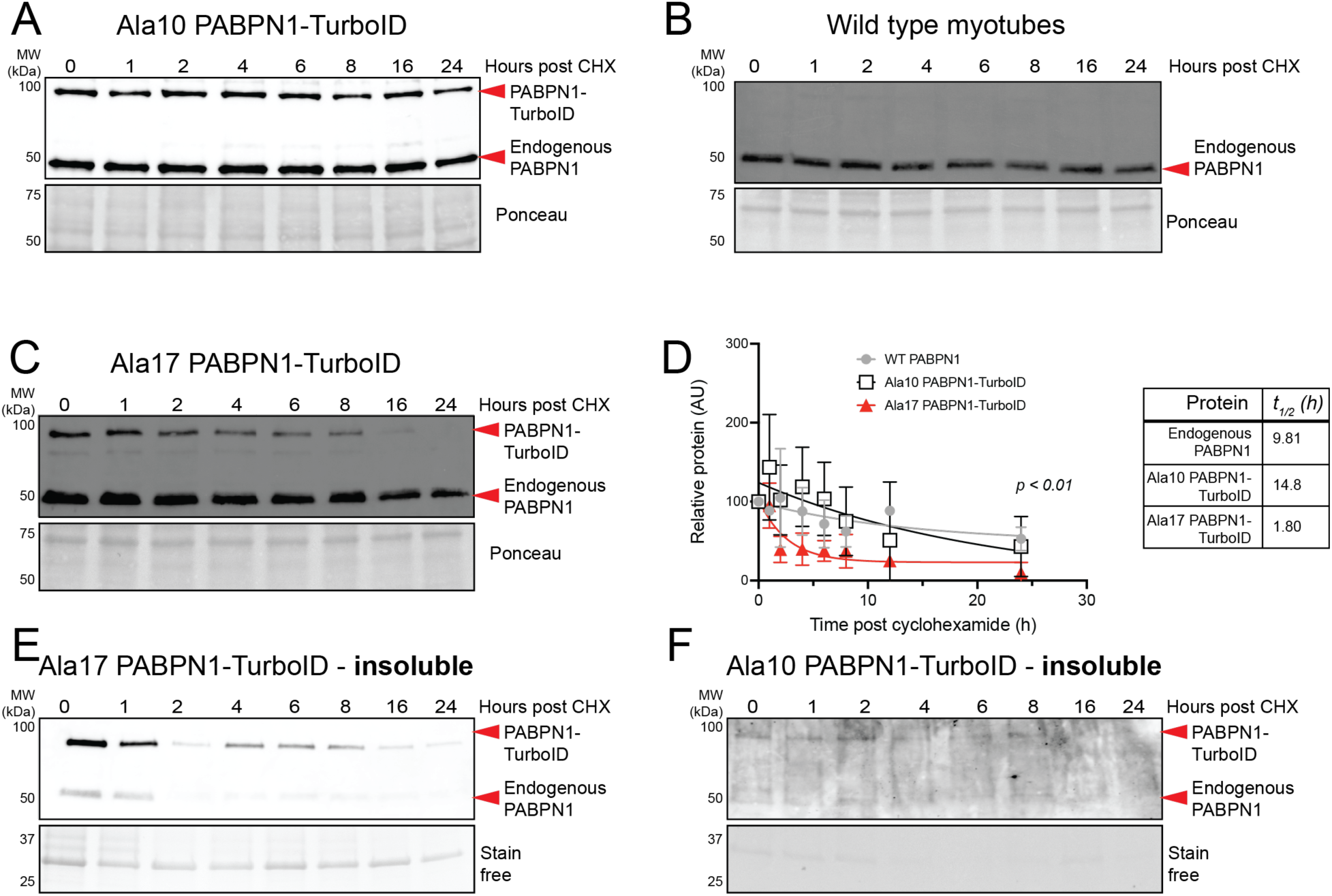
Reduced half-life of soluble and insoluble Ala17 PABPN1-TurboID compared to Ala10 PABPN1-TurboID and endogenous PABPN1. **A)** Immunoblot of cycloheximide (CHX) chase experiments showing similar stability of Ala10 PABPN1-TurboID and endogenous PABPN1 as detected by an antibody to PABPN1 in myotubes expressing near native levels of Ala10 PABPN1-TurboID. **B)** Immunoblot of cycloheximide chase experiments showing stability of endogenous PABPN1 in wild type cells. **C)** Immunoblot of cycloheximide chase experiments showing relative instability of Ala17 PABPN1-TurboID in myotubes expressing near native levels of Ala17 PABPN1-TurboID with little effect on endogenous PABPN1 stability using an antibody to PABPN1. For *A-C*, Ponceau stain was used as a loading control. **D)** Non-linear fit and one-phase decay calculation showing reduced half-life of Ala17 PABPN1-TurboID relative to Ala10 PABPN1-TurboID and endogenous PABPN1. Shown is mean ± standard deviation for n = 3-5 experiments. Statistical significance was determined by mixed-effects analysis with Tukey’s post-hoc test. p < 0.05 for comparison of Ala10 PABPN1-TurboID to endogenous PABPN1. p < 0.01 for comparison between Ala17 PABPN1-TurboID and endogenous PABPN1. p < 0.0001 for comparison between Ala17 and Ala10 PABPN1-TurboID. **E)** Immunoblot of insoluble fractions from cycloheximide chase experiment showing presence of Ala17 PABPN1-TurboID and endogenous PABPN1 in myotubes expressing Ala17 PABPN1-TurboID as detected using an antibody to PABPN1. Reduced stability of both Ala17 PABPN1-TurboID and endogenous PABPN1 can be observed relative to PABPN1 in the soluble fractions shown in *A-C*. **F)** Immunoblot of insoluble fractions from cycloheximide chase experiment showing little to no Ala10 PABPN1-TurboID or endogenous PABPN1 in myotubes expressing Ala10 PABPN1-TurboID. For *E* and *F*, stain-free gel imaging used as a loading control.

Single-phase decay calculations revealed that the half-life of endogenous PABPN1 in wild type C2C12 myotubes was ∼10h while the half-lives of the Ala10 and Ala17 PABPN1-TurboID constructs were ∼15h and ∼2h, respectively (Figure 5D). These data indicate that soluble Ala17 PABPN1-TurboID is significantly less stable than Ala10 PABPN1-TurboID or endogenous wild type PABPN1. A decrease in Ala17 PABPN1-TurboID protein over time could be a result of increased movement of PABPN1 into the insoluble fraction. To detect insoluble PABPN1, insoluble pellets from cells used for cycloheximide experiments were solubilized in urea and analyzed by immunoblot. Both Ala17 PABPN1-TurboID and endogenous PABPN1 were detected in insoluble pellets from Ala17 PABPN1-TurboID expressing myotubes and the levels of both proteins decreased similarly to soluble protein after cycloheximide treatment (Figure 5E). In myotubes expressing Ala10 PABPN1-TurboID, very little insoluble PABPN1 was detected (Figure 5F). These results suggest that alanine expansion leads to increased degradation and decreased solubility of PABPN1, which may impact PABPN1 function.

Beyond its canonical function of stimulating poly(A) polymerase, PABPN1 promotes nuclear export of mature, polyadenylated RNAs which is mediated by the transcription export (TREX) complex^38^. Major components of the TREX complex including Aly/REF, THOC1, and THOC5^39,40^ were not detected in any of the PABPN1 proximal proteins identified by proteomics, which may be a result of their relatively low expression in myotubes. Immunoblot confirmed that levels of Aly/REF, THOC1, and THOC5 were significantly reduced in myotubes compared to undifferentiated myoblasts (Figure 6A, B). Increased detection of Ala17 PABPN1 proximal proteins related to splicing and polyadenylation and decreased detection of the cytosol facing nuclear pore protein NUP214 compared to Ala10 PABPN1 proximal proteins in myotubes suggests that alanine expansion might affect PABPN1 interaction with nuclear export machinery. Indeed, significantly lower levels of THOC1 were detected in immunoblots of streptavidin eluates from Ala17 PABPN1-TurboID expressing myotubes relative to those expressing Ala10 PABPN1-TurboID while detection of Aly/REF and THOC5 trended downward (Figure 6C, D). Decreased labeling of nuclear export machinery by Ala17 PABPN1-TurboID suggests that Ala17 PABPN1 may be less efficient than Ala10 PABPN1 at promoting export of polyadenylated mRNAs.

**Figure 6:**
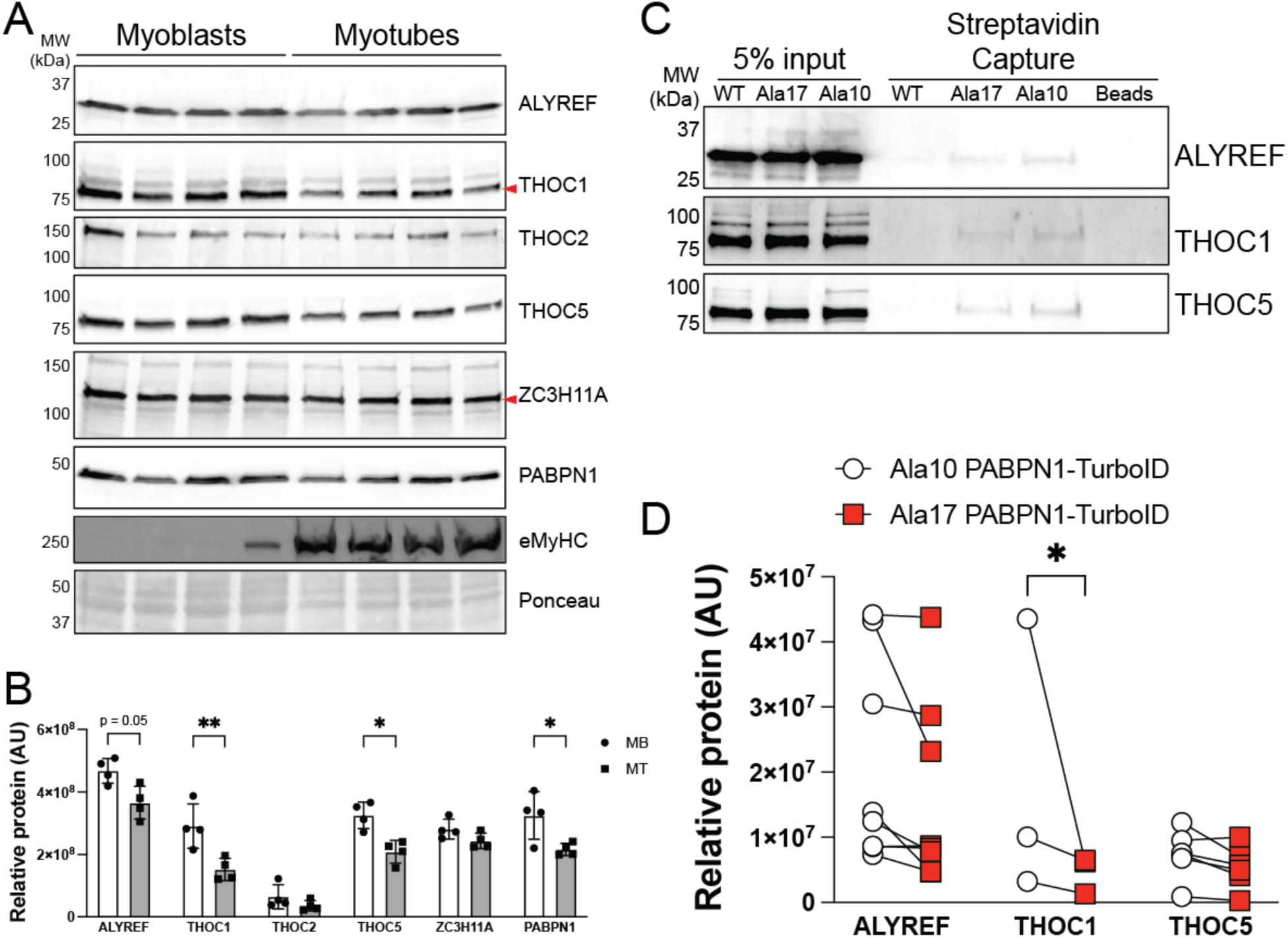
Components of the RNA export machinery pathway decrease in abundance during myoblast differentiation and are labeled less by Ala17 PABPN1-TurboID. **A)** Immunoblots probed with antibodies to various components of TREX complex (ALYREF, THOC1, THOC2, THOC5) comparing levels in myotubes compared to myoblasts. Blots were probed with antibody to PABPN1 as a positive control for reduced levels of RNA processing machinery in myotubes and an antibody to myosin heavy chain (eMyHC) as a marker of differentiation. Ponceau stain was used as a loading control. **B)** Quantification of blot shown in *A* showing significantly reduced levels of THOC1, THOC5, and PABPN1 in myotubes compared to myoblasts. Shown is mean ± standard deviation for n = 4 experiments. **C)** Immunoblot of streptavidin elutions from myotubes expressing Ala10 and Ala17 PABPN1-TurboID probed with antibodies to ALYREF, THOC1, and THOC5. **D)** Quantification of experiments represented in *C*. Shown is mean ± standard deviation for n = 3-7 experiments. For B and D, statistical significance was determined using two-way ANOVA with Šídák’s multiple comparisons test. * p < 0.05, ** p < 0.01

To determine the effect of Ala17 PABPN1 expression on nuclear poly(A) RNA export, oligo d(T) FISH was performed in myotubes expressing near-native levels of Ala10 or Ala17 PABPN1-TurboID. Wild type myotubes and those expressing Ala10 PABPN1-TurboID showed similar distribution of polyadenylated RNAs in both the nucleus and cytosol (Figure 7A). However, myotubes expressing Ala17 PABPN1-TurboID showed significant nuclear accumulation of polyadenylated RNAs that could be detected in a punctate pattern (Figure 7A, B). Overexpression of Ala17 PABPN1-TurboID also led to a significant increase in nuclear accumulation of polyadenylated RNA and nuclear RNA puncta could be detected in cells overexpressing either Ala10 or Ala17 PABPN1-TurboID (Figure 7C, D), though this effect was more with Ala17 PABPN1-TurboID overexpression. To confirm these results, we performed oligo d(T) FISH in primary myotubes generated from the *Pabpn1^+/A17^* knock-in mouse model of OPMD^21,29^. Like C2C12 myotubes expressing Ala17 PABPN1-TurboID, reduced poly(A) RNA signal in the cytoplasm could be observed in myotubes from *Pabpn1^+/Ala17^* mice (Figure 7E). Considering that Ala17 PABPN1-TurboID expressing myotubes and primary *Pabpn1^+/Ala17^* myotubes contain at least one endogenous *Pabpn1* allele, these data indicate that alanine expansion elicits a dominant negative effect on PABPN1-dependent nuclear export of polyadenylated RNA.

**Figure 7:**
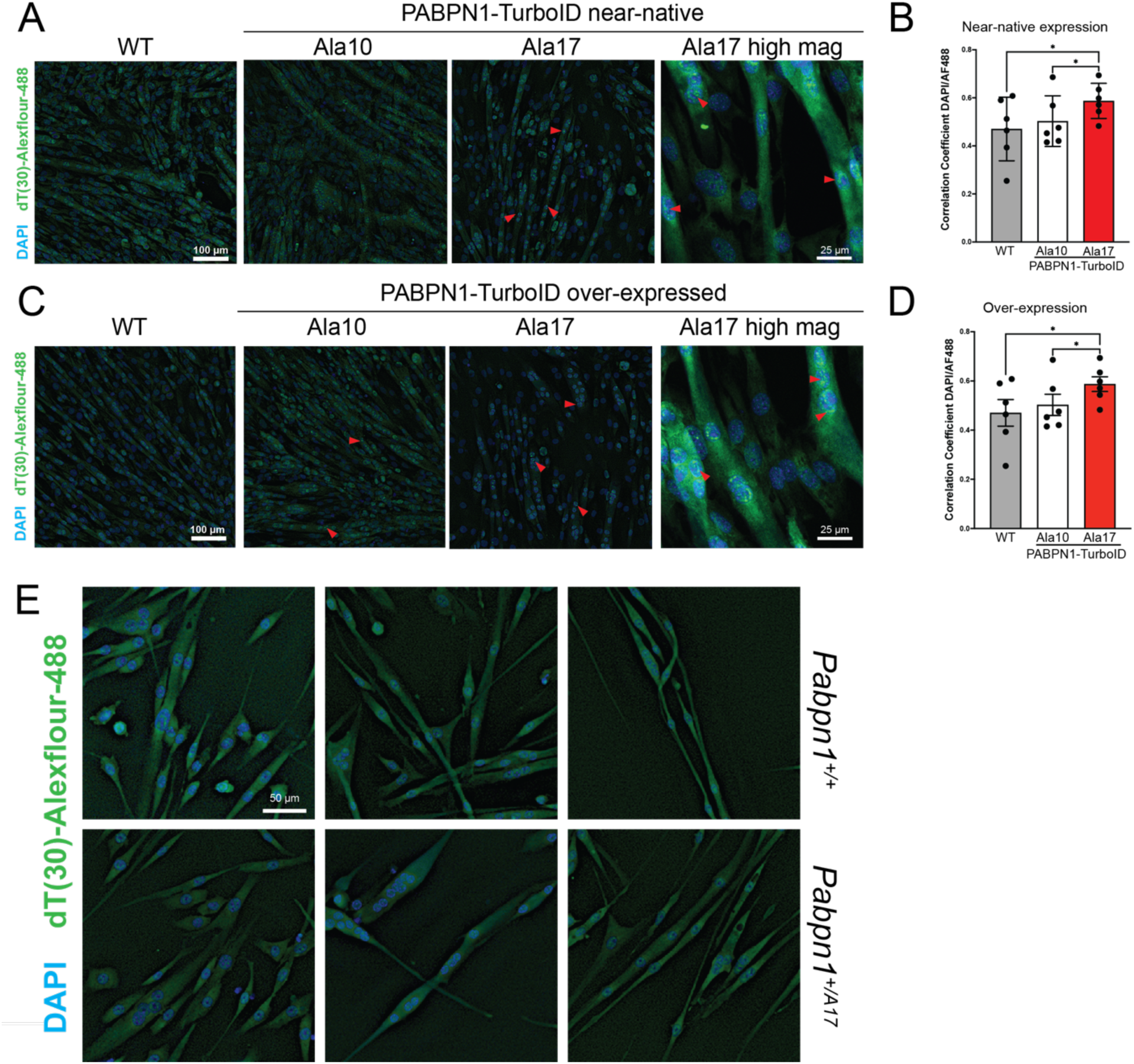
Ala17 PABPN1 causes a dominant negative effect on polyadenylated RNA localization. **A)** Fluorescence in situ hybridization (FISH) in wild type (WT) myotubes or myotubes expressing near-native levels of Ala10 or Ala17 PABPN1-TurboID using d(T)_30_-AF-488 to detect polyadenylated RNA and DAPI to stain nuclei. Additional high magnification images of Ala17 PABPN1-TurboID expressing myotubes shown on the right. Red arrows mark polyadenylated RNA puncta. **B)** Quantification of Pearson’s correlation showing increased co-localization of DAPI-stained nuclei with d(T)_30_-AF-488 in myotubes expressing Ala17 PABPN1-TurboID relative to those expressing Ala10 PABPN1-TurboID at near-native levels or WT myotubes. **C)** d(T)_30_ FISH in WT myotubes or myotubes overexpressing Ala10 or Al7 PABPN1-TurboID. Additional high magnification images of Ala17 PABPN1-TurboID expressing myotubes shown on the right. Red arrows highlight polyadenylated RNA puncta. **D)** Quantification of Pearson’s correlation showing increased co-localization of DAPI-stained nuclei with d(T)_30_-AF-488 in myotubes over-expressing Ala17 PABPN1-TurboID relative to those over-expressing Ala10 PABPN1-TurboID or to the WT myotubes quantified in *B*. For *B* and *D*, shown is mean ± standard deviation for n = 6 experiments. Statistical significance determined using one-way ANOVA. * p < 0.05. **E)** d(T)_30_-AF-488 FISH showing reduced polyadenylated RNA signal in the cytoplasm of primary myotubes from a knock-in mouse model of OPMD (*Pabpn1^+/A17^*, bottom) compared to wild-type litter mates (*Pabpn1^+/+^*, top). Shown is one representative image each from three different mice per genotype.

## Discussion

Here, we focused on understanding the effect of OPMD-associated alanine expansion on PABPN1 function. Stable C2C12 myoblast lines expressing Ala10 or Ala17 PABPN1-TurboID constructs under the control of an inducible promoter allowed for comparison of function and protein binding partners without the confounding effects of overexpression. Experiments to test the functionality of PABPN1-TurboID constructs revealed that, while Ala10 PABPN1-TurboID was able to completely complement endogenous PABPN1 deficiency, Ala17 PABPN1-TurboID could not. Additionally, expression of Ala17 PABPN1-TurboID impaired some PABPN1 functions even in the presence of endogenous PABPN1. Finally, comparative proteomic analysis showed that, while both Ala10 and Ala17 PABPN1 are found in close proximity to similar cohorts of proteins, many of those involved in splicing, cleavage and polyadenylation, and protein turnover were enriched in Ala17 compared to Ala10 PABPN1-TurboID proximal proteins. Thus, data generated from these PABPN1-TurboID constructs support a model where alanine expansion both impairs PABPN1 function and causes a dominant negative effect.

The continuum of possible molecular mechanisms underlying OPMD pathology proposed by Banerjee et al includes the potential for PABPN1 function to be directly affected by alanine expansion^1^. Here, we show that the RNA export or differentiation defect caused by knocking down endogenous PABPN1 can be restored to wild type levels by expressing Ala10 but not Ala17 PABPN1-TurboID. However, the cell death phenotype caused by *Pabpn1* deficiency under differentiation conditions was rescued by both Ala10 and Ala17 PABPN1-TurboID. These data suggest that Ala17 PABPN1 is less effective than Ala10 PABPN1 at mediating some but not all PABPN1 functions. Several other lines of evidence point to a loss of PABPN1 function caused by alanine expansion. Our previous studies reported reduced poly(A) tail length in both the *Pabpn1^+/Ala17^* knock-in mouse model of OPMD and in a hemizygous PABPN1 knockout (*Pabpn1^+/Δ^*) mouse, suggesting that the canonical PABPN1 function of promoting poly(A) polymerase activity is impaired by alanine expansion^21^. However, mitochondrial dysfunction observed in muscles from *Pabpn1^+/Ala17^* mice were not observed in *Pabpn1^+/Δ^* mice. A recent study comparing sequencing data from a cell culture model of OPMD and muscle tissue from OPMD patients suggested that shifts in alternative polyadenylation signal (PAS) usage correlated with loss of alternative isoforms of the *PABPN1* transcript^41^. Finally, a study exploring the efficacy of a knockdown and replace gene therapy for OPMD^22^ compared OPMD-associated phenotypes in the A17.1 mouse model and in primary myoblasts isolated from an individual with OPMD. For several phenotypes tested, including muscle fibrosis, myofiber cross-sectional area, and cell death, vector-encoded Ala10 PABPN1 was sufficient to reverse the phenotypes caused by alanine-expanded PABPN1. These results showed that some pathologic hallmarks of OPMD may be driven by loss of PABPN1 function. However, some functional and molecular phenotypes were not resolved by adding back Ala10 PABPN1 alone but instead required simultaneous knockdown of expanded PABPN1. The partial rescue of OPMD-associated phenotypes by Ala10 PABPN1 suggests that alanine expansion imparts additional dominant negative phenotypes.

Indeed, data reported here support a model wherein alanine expansion exerts a dominant negative effect on PABPN1 function even in the absence of overexpression. Ala17 PABPN1-TurboID elicited a mild differentiation defect in control cells without *Pabpn1* knockdown. These results are consistent with the in vivo and in vitro differentiation defects we previously reported in pharynx-derived satellite cells and myoblasts in the *Pabpn1^+/A17^*knock-in mouse model of OPMD^29^. Additionally, near native levels of Ala17 PABPN1-TurboID caused nuclear accumulation of polyadenylated RNA in C2C12 and primary myotubes as detected by oligo d(T) FISH experiments even in the presence of endogenous wild type PABPN1. Interestingly, polyadenylated RNA formed puncta in the nuclei of C2C12 myotubes expressing near native levels of Ala17 but not Ala10 PABPN1-TurboID. These puncta were larger and more visible upon overexpression where even the presence of Ala10 PABPN1-TurboID led to some observable punctate oligo d(T) staining. Surprisingly, though insoluble PABPN1 was detected in cells expressing Ala17 PABPN1-TurboID, no nuclear PABPN1 aggregates were detected in myotubes immunostained with an antibody to PABPN1. Though PABPN1 aggregates are a pathologic hallmark of OPMD, they are only detected in 5-10% of myonuclei in limb muscle biopsies from affected individuals^18^ and the knock-in mouse model of OPMD^21^. Interestingly, a recent study by Roth et al.^18^ showed reduced numbers of PABPN1 aggregates in cricopharyngeal muscles compared to other muscles, which exhibit higher nuclear turnover than limb muscles and higher basal activity of muscle stem cells^29,42,43^. PABPN1 aggregates were also not detected in OPMD patient-derived xenografts in mouse muscles, formation of which requires a step of degeneration and regeneration. The authors of this study hypothesized that expanded PABPN1 does not form detectable aggregates in newly fused myonuclei and that PABPN1 aggregates only appear in aged nuclei. Our data support this model and the lack of PABPN1 aggregates detected in Ala17 PABPN1-TurboID expressing myotubes may be due to the fact that expanded PABPN1 is expressed for only 48 hours in this system. The insoluble PABPN1 and polyadenylated RNA puncta we detected in Ala17 PABPN1-TurboID expressing myotubes may represent stalled RNA processing complexes that later develop into aggregates, which have previously been described as pre-intranuclear inclusions (INI)^44^. Thus, our data and the study by Roth et al suggest a model in which short-term expression of expanded PABPN1 is not sufficient to promote aggregate formation.

Comparative proteomics revealed that components of the splicing and polyadenylation machinery were enriched in Ala17 versus Ala10 PABPN1 proximal proteins. This result would be consistent with stalled RNA processing caused by alanine expansion of PABPN1. Indeed, previous studies using glycerol gradient centrifugation found that PABPN1 is detected in higher molecular weight complexes, and these complexes are larger in myotubes expressing alanine-expanded PABPN1^24,35^. Future studies will be needed to probe the effect of alanine expanded PABPN1 on the kinetics of RNA processing complex assembly, activity, and disassembly. Proximal proteins that were detected equally in both Ala10 and Ala17 PABPN1-TurboID data sets were enriched in GO terms/pathways related to ER stress. Previous studies have suggested that expression of alanine-expanded PABPN1 can induce ER stress and the unfolded protein response (UPR) potentiator guanabenz was reported to reduce ER stress and ameliorate OPMD phenotypes in the A17.1 mouse model^45^. It is possible that ER stress-related proteins are detected due to heterologous expression of exogenous PABPN1 in the A17.1 mouse model of OPMD and in PABPN1-TurboID expressing myotubes reported here. However, it may be the case that both Ala10 and Ala17 PABPN1 interact with components of the ER stress machinery but alanine expansion of PABPN1 aberrantly affects this interaction leading to ER stress. Finally, we detected multiple overlapping proteins and pathways detected in our previous study of immunoprecipitated tagged PABPN1 in mouse skeletal muscle^24,35^. However, many of these proteins, like those defined as mitochondria-localized, were not detected as PABPN1-TurboID proximal proteins in this study. The absence of mitochondrial proteins detected by PABPN1-TurboID could indicate that PABPN1 precipitation of mitochondrial proteins represents a non-specific interaction generated by overexpression of an exogenous tagged protein or could be related to the use of C2C12 myotubes rather than murine skeletal muscle in this study.

The use of C2C12 myotubes is indeed a limitation of this work. For example, some known PABPN1 functions, like its role in promoting DNA double strand break (DSB) repair^46^ were not represented in the PABPN1 proximal protein data generated here. DSB is active in satellite cells but declines during myoblast differentiation^47,48^ so the effect of alanine expansion on PABPN1 function may be detectable in proliferating myoblasts. Similarly, phosphorylated PABPN1 was recently reported promote transcript stability and proliferation in HeLa and HEK293 cells^49^. We have previously demonstrated that basal autophagy and pharynx-derived myoblast proliferation is impaired in primary cells isolated from the *Pabpn1^+/A17^* knock-in mouse model of OPMD^29^. The effect of alanine expansion on PABPN1 post-translational modification and function is likely to differ by both differentiation state and specific muscle or muscle group. Future studies focusing on dissecting the effect of alanine expansion on PABPN1 function in a differentiation and muscle specific context will help reveal why OPMD primarily affects specific muscle groups.

## Materials and Methods

### Plasmids and cloning

Sequences encoding Ala10 PABPN1-TurboID or Ala17 PABPN1-Turbo constructs with two glycine serine hinges, a DYKDDDDK (Flag) tag, TurboID, and a Myc tag fused onto the C-terminus of PABPN1 were synthesized by Azenta. This expression cassette was preceded by a strong Kozak sequence. The expression cassette was excised from the puc57 shuttle vector using HindIII and XbaI and inserted into a pcDNA3.1 backbone. Plasmids were confirmed via Sanger sequencing. To generate stable cells, expression cassettes encoding either Ala10 or Ala17 PABPN1-TurboID were cloned into the pCW57.1 lentiviral vector (pCW57.1 was purchased from Addgene after being donated by David Root; Addgene plasmid # 41393; http://n2t.net/addgene:41393; RRID:Addgene_41393) containing a doxycycline inducible promoter and puromycin resistance selectable marker using EcoR1 restriction sites added via PCR. Plasmids were confirmed by Sanger sequencing.

### Cell culture

C2C12 myoblasts were purchased from ATCC (CRL-1772) and HEK 293T cells were a kind gift from Dr. William Miller (University of Cincinnati). Growth medium for C2C12 and HEK 293T cells included Dulbecco’s Modified Eagle Medium (DMEM) with 4.5 g/L glucose, L-glutamine, and sodium pyruvate (Corning 10-013-CV) plus 10% fetal bovine serum (FBS, Cytiva SH3091003) and 100 µg/ml penicillin-streptomycin (Corning 30-002-CI) with a 1:10,000 maintenance dose of the anti-mycoplasma reagent Plasmocin (Invivogen ant-mpt-1). To differentiate C2C12 myoblasts into mature myotubes, confluent cells were provided with differentiation medium containing DMEM with 2% Horse Serum (Cytiva SH3007403) and 100 µg/ml penicillin-streptomycin for 72-120 hours. Differentiation medium was refreshed approximately every 48 hours. Complete differentiation was determined by morphology, confirmed by expression of embryonic myosin heavy chain, and quantified by fusion index (% nuclei in multinucleated myotubes relative to total nuclei). Fusion index was calculated in a blinded fashion using images chosen randomly from a minimum of three areas per well.

Stocks of primary myoblasts isolated from the knock-in mouse model of OPMD (*Pabpn1^+/A17^*) or wild type littermates (*Pabpn1^+/+^*) used in Zhang et al.^29^ were used to confirm d(T)_30_-AF488 FISH results. Cells were grown in dishes pre-coated with bovine collagen I (Corning 354231) in HyClone Ham’s F10 medium (Cytiva SH3002501) with 20% FBS, 100 µg/ml penicillin-streptomycin, and 5ng/mL human basic fibroblast growth factor (Pepro Tech 100-18B). To differentiate primary myoblasts, cells were seeded on plates were pre-coated with entactin-collagen IV-laminin (Sigma 08-110) and provided with DMEM containing 1 g/L glucose, sodium pyruvate, and L-glutamine (Gibco 11885084) with 1x P/S and 1% insulin-selenium-transferrin solution (Gibco 51300044). Differentiation took 72 hours and was assessed by morphology.

### Generation of stable cell lines

HEK 293T cells (3.5 x 10^5^ cells/well) were seeded onto six well plates pre-coated with Matrigel (Corning 356234) and allowed to attach overnight. Cells were transfected with plasmids encoding either Ala10 or Ala17 PABPN1-TurboID alongside plasmids PsPax2, a packaging construct containing HIV GAG-Pol, and PMDG.2, a viral envelope expressing plasmid containing VSVG Envelope. These plasmids were transfected at a ratio of 4:2:1 using Lipofectamine 3000 (Invitrogen L3000001). After approximately 16 hours, transfection medium (growth medium without P/S) was removed, and growth medium was added to each well. To select transfected cells, 2 µg/ml puromycin (Gibco A1113803) was added to each well.

Approximately 48-72 hours later, enriched medium was removed from transfected HEK293T cells and centrifuged twice at 500 x g (5 minutes) to remove any remaining cells and further clarified by passing through a Whatman PES 0.45 µm syringe filter. Enriched medium containing lentiviral particles (3 ml) was diluted into growth medium (7 ml) containing 8 µg/ml (final) polybrene (Sigma TR-1003). Remaining vector was frozen overnight at -80 °C and then moved to liquid nitrogen storage. Vector-containing enriched medium was added to 10 cm plates previously seeded with low passage C2C12 myoblasts. To ensure even distribution of vector, plates that received enriched medium were rocked back and forth repeatedly every 30 minutes for approximately 8 hours. After 24 hours, transduced C2C12 myoblasts were selected by switching to fresh growth medium containing 2 µg/ml puromycin. Selection was performed again 48 hours after transduction and continued approximately every other passage for the duration of cell growth.

### Induced expression of PABPN1-TurboID and labeling of proximal proteins

Wild type and transduced C2C12 myoblasts containing either Ala10 or Ala17 PABPN1-TurboID were grown to 90-95% confluency and then induced to differentiate. Upon observation of mature myotubes, differentiation medium was removed and fresh medium containing 1-2 µg/ml doxycycline was added to induce expression of Ala10 or Ala17 PABPN1-TurboID. For dose response experiments the concentration of doxycycline ranged from 1-10 µg/ml. Medium containing doxycycline was refreshed 24 hours later. Approximately 48 hours after doxycycline was added, myotubes were treated with 1 mM biotin for two hours or harvested as needed.

### Immunoblotting

Prior to harvest, cells were washed twice in ice-cold phosphate buffered saline (PBS, 137 mM NaCl, 2.7 mM KCl, 11.9 mM phosphates pH 7.4, purchased as 10X solution packets, Fisher BioReagents BB6651). Lysates were then collected in either 1x RIPA buffer (Pierce PI89900) containing 1x protease inhibitor (Pierce PIA32953) and 1% SDS or harsh lysis buffer (8M urea and 2% SDS). Lysates were processed by sonicating for ten seconds twice each at low power with one minute rest on ice. To avoid excess precipitation of aggregate-prone PABPN1 we did not clarify via high-speed centrifugation for most experiments. For cycloheximide experiments, protein lysates were clarified by high-speed centrifugation for 30 minutes at 4 °C. Upon collection, processing, and clarification of protein lysates, insoluble pellets were washed once in 1x PBS, resuspended in a solubilization buffer containing 8 M urea and 2% SDS, and boiled for 5 minutes. Lysate concentrations were quantified by Bradford assay and 30 ug of protein separated on 4-20% acrylamide Mini-Protean TGX Stain-Free PAGE gels (BioRad 4568094).

Gels were either stained with silver stain (Pierce 24612) according to the manufacturer’s instructions or transferred to a nitrocellulose membrane. Stain-free imaging technology (BioRad) or Ponceau stain was used to assess transfer and loading for downstream quantification. Blots were blocked for 15 minutes to 1 hour in Everyblot solution (BioRad 12010020) and probed overnight at 4 °C in primary antibody diluted in Tris buffered saline (20 mM Tris, 150 mM HCl pH 7.6) with 0.1% Tween-20 (TBST) followed by 1 hour at room temperature in secondary antibody diluted in 5% milk and TBST. A complete list of antibodies and dilutions can be found in Supplementary Table 1. Immunoblots were developed using ECL Supersignal or Femto plus (Pierce PI34580 or PI34095) and visualized on a ChemiDoc imager (BioRad). Immunoblot images were quantified by densitometry using Image Lab software (BioRad).

### Subcellular fractionation

To isolate the nuclear fraction of C2C12 myotubes, a modified version of the nuclear fractionation protocol described by Bannerjee et al.^35^ was used. Cells were scraped into ice cold PBS and centrifuged at 500 x g for 5 minutes at 4 °C. Cell pellets were resuspended in NP-40 lysis buffer (10 mM Tris-HCl, pH 7.4, 10 mM NaCl, 3 mM MgCl_2_, 1% [v/v] igepal detergent) with gentle pipetting, incubated on ice for 5 minutes, gently flicking the tube every minute. 40 µl of total cell lysate was removed, and the remaining lysate was centrifuged at 500 x g at 4 °C for 5 minutes. Supernatant was removed as the cytoplasmic fraction. The nuclear pellet was washed in PBS, centrifuged, and the pellet resuspended in RIPA buffer with 1% SDS. Sub-cellular fractionation results were evaluated via immunoblot, probing for PABPN1, GAPDH as a marker of the cytoplasmic fraction, and Histone H3 as a marker of the nuclear fraction.

### Pabpn1 knockdown and rescue with PABPN1-TurboID constructs

PABPN1 deficiency was induced by siRNA as described^29^. C2C12 myoblasts were transfected with an siRNA specific to the *Pabpn1* 3’ untranslated region (UTR). The *Pabpn1* 3’ UTR targeting siRNA did not successfully reduce endogenous PABPN1 protein during differentiation, so a coding sequence (CDS) targeting siRNA was used. Both siRNAs were purchased as dicer substrate siRNAs from IDT (*Pabpn1* 1.13.3 for 3’UTR targeting and *Pabpn1* 1.13.1 for CDS targeting). Myoblasts were provided with *Pabpn1* targeting or non-targeting control siRNAs packaged in Lipofectamine 3000 for 12-16 hours in growth medium without P/S and then switched to fresh medium. For experiments in myoblasts, cells were switched to normal growth medium for an additional 48 hours before harvest. For experiments across differentiation, cells were provided with differentiation medium and harvested when control wells showed complete myotube formation. For rescue experiments in myoblasts, stable transduced cells were treated with 2 µg/ml doxycycline when switched to fresh medium 12-16 hours after transfection. Addition of doxycycline affected differentiation of control cells, so cells induced to differentiate were instead co-transfected with pcDNA 3.1 plasmid encoding either Ala10 or Al17 PABPN1-TurboID. *Pabpn1* knockdown and TurboID construct re-expression was confirmed by immunoblot.

### Immunofluorescence staining and microscopy

C2C12 myoblasts or myotubes were fixed using 3.7% paraformaldehyde (Alfa Aesar AA433689M) in PBS for 10 minutes, washed 3 times in PBS for 5 minutes, and permeabilized using PBS containing 0.3% TritonX-100 for 15 minutes while rocking. Blocking buffer containing 3% BSA in 1x PBS with 0.3% TritonX-100 was added to the plates for 1 hour at room temperature. Primary antibodies (Supplementary Table 1) resuspended in 0.5x blocking buffer were added to each well and incubated overnight 4 °C. The next day each plate was washed and secondary antibody mixtures suspended in 0.5x blocking buffer added to the cells for 1 hour at room temperature while rocking. Nuclei were stained with 4^’^,6-diamidino-2-phenylindole (DAPI) for 10 minutes at room temperature. Staining was visualized and images recorded using an Olympus IX83 fluorescence microscope with an Olympus DP74 camera using by CellSens Dimensions imaging software. Images were taken from at least three areas per well chosen randomly. Images were subjected to downstream analysis in FIJI^50^.

### Oligo d(T) fluorescence in situ hybridization (FISH), imaging, and quantification

C2C12 myoblasts or myotubes and primary myotubes were washed 3 times in ice-cold RNA safe PBS containing 0.1% diethyl pyrocarbonate (DEPC, Research Products International 50-213-289) and RNAse inhibitor (Promega RNasin PRN2611) for five minutes and fixed with 3.7% paraformaldehyde in RNA safe PBS for 10 minutes at room temperature. Cells were then washed with RNA safe PBS 3 times for five minutes, and permeabilized with ice cold methanol containing RNasin for 10-15 minutes on ice while rocking. Additional permeabilization occurred overnight at 4 °C in 70% ethanol with RNasin. The following day, cells were hybridized with 0.5 µM d(T)_30_ conjugated to AlexaFluor-488 (IDT) for 1 hour at 37 °C in 0.2 mM hybridization buffer (25% formamide, 10% dextran sulfate, 2x SSC [3 M sodium chloride and 300 mM trisodium citrate at pH 7.0]) with RNasin and protected from light. Cells were washed twice for 5 minutes each with 2x SSC buffer, washed 3x with RNA safe PBS, and nuclei were stained with DAPI. Stained cells were visualized on an Olympus IX83 fluorescence microscope as described above or on a Leica Stellaris 8 confocal microscope (University of Cincinnati College of Medicine Live Microscopy Core). Pearson’s correlation of AF-488 and DAPI signal as calculated by CellProfiler was used to quantify colocalization of polyadenylated RNA and nuclei as described in the included example protocol^51^.

### Streptavidin precipitation

Lysates were harvested in 1x RIPA buffer containing 1x protease inhibitor and 1% SDS. Lysates were sonicated twice for 10 seconds each on ice. Streptavidin conjugated agarose beads (Pierce 20357) were pre-cleared using 1% bovine serum albumin (BSA) in IP buffer (50 mM Tris-HCl pH 7.4, 150 mM NaCl, 0.5% Igepal). Beads were washed gently 3-4 times with IP buffer to remove any residual BSA, 1-2 mg of lysate was added to the beads, and diluted to a volume of 1 ml with IP buffer. Bead-lysate mixtures were allowed to tumble end-over-end overnight at 4 °C. Beads were centrifuged at 800 x g and residual lysate aspirated. Beads were then washed 5 times with 1000 µl of IP buffer each. Precipitated biotinylated protein was eluted by adding 200 µl of 8M urea, 2% SDS, and 40 µl of Laemmli buffer and boiling for five minutes. Samples were analyzed by silver staining to assess yield, blots probed with specific antibodies or streptavidin conjugated to HRP to assess labeling, and proteomics to identify total proximal proteins.

### Comparative Proteomics

Three separate experiments were performed to elute biotinylated proteins from wild type myotubes or myotubes expressing near-native levels of Ala10 or Ala17 PABPN1-TurboID treated with doxycycline to induce expression and biotin as a substrate for the TurboID. Comparative proteomics were performed at the University of Cincinnati Proteomics Laboratory (Core Director: KD Greis, Ph.D.) on biotinylated proteins enriched by streptavidin capture and eluted in Laemmli sample buffer with urea (described above) using label-free quantitative nanoLC-MS/MS workflow as previously described^52^. Briefly, samples were run ∼ 2cm into a 4-12% Bis-Tris gel (Invitrogen) using MOPS buffer and separated by pre-stained molecular weight markers. All proteins in the gel were excised, reduced with dithiothreitol, alkylated with iodoacetic acid, and digested overnight with trypsin. Resulting peptides were analyzed by nanoLC-MS/MS on a Dionex Ultimate 3000 RSLCnano coupled to a Thermo Orbitrap Eclipse mass spectrometry system. Data were collected using Xcaliber 4.3 software (Thermo Scientific) with label-free quantitation and comparative profiling of proteins detected from each sample group performed using Proteome Discoverer 3.0 (Thermo Scientific) against the complete *Mus musculus* protein database. Proteins with the minimum of 2 high (99%) confidence peptides were reported. In comparing between groups, strict analysis included proteins detected in two of three lysates generated from PABPN1-TurboID expressing cells but not detected in wild type controls. A more inclusive analysis was also performed on PABPN1-TurboID proximal proteins that were detected at least 2-fold higher than those detected in wild type control samples. Functional annotation was performed using STRING database version 12.0^30^.

### Cycloheximide chase assay

PABPN1 and PABPN1-TurboID protein stability was calculated by cycloheximide chase assay. Approximately 48 hours after expression was induced, wild type myotubes or myotubes expressing Ala10 or Ala17 PABPN1-TurboID were incubated with 50 µg/ml cycloheximide (ThermoFisher J66004). Lysates were harvested in RIPA at 0, 1, 2, 4, 6, 8, 12, and 24 hours and analyzed by immunoblot. All samples were clarified after sonication and insoluble pellets were retained and resuspended in harsh lysis buffer as described above. Protein levels were quantified by densitometry using Image Lab software (BioRad). PABPN1 half-life was calculated using non-linear fit and one-phase decay analysis in GraphPad Prism 10 for macOS.

### Statistical analysis

Statistical analysis was performed using GraphPad Prism 10 for macOS. The statistical calculations used, sample size, and p value range for each experiment is included in the figure legend. For most experiments, shown are individual values and mean ± standard deviation. Figures 3I, 6D, and S4C show individual values paired with controls. For protein half-life calculations shown in Figure 5D, non-linear fit shows mean ± standard deviation without individual values. For experiments with one variable, two groups were compared using paired t-test unless otherwise noted and three or more groups were compared using one-way ANOVA with Dunnett’s post-hoc correction for multiple comparisons. For experiments comparing more than one related variable and two or more groups, two-way ANOVA with Tukey’s post hoc correction for multiple comparisons was used as stated in the figure legend. In all cases, p < 0.05 was considered to be statistically significant.

## Acknowledgements

The authors would like to acknowledge the University of Cincinnati core facilities employed to generate data reported here. Mass spectrometry data were collected and analyzed in the University of Cincinnati College of Medicine Proteomics Laboratory under the direction of KD Greis, Ph.D. Confocal microscopy was performed in the University of Cincinnati College of Medicine Live Microscopy Core.

## Figures and Supplemental Figures

**Supplemental Figure 1:**
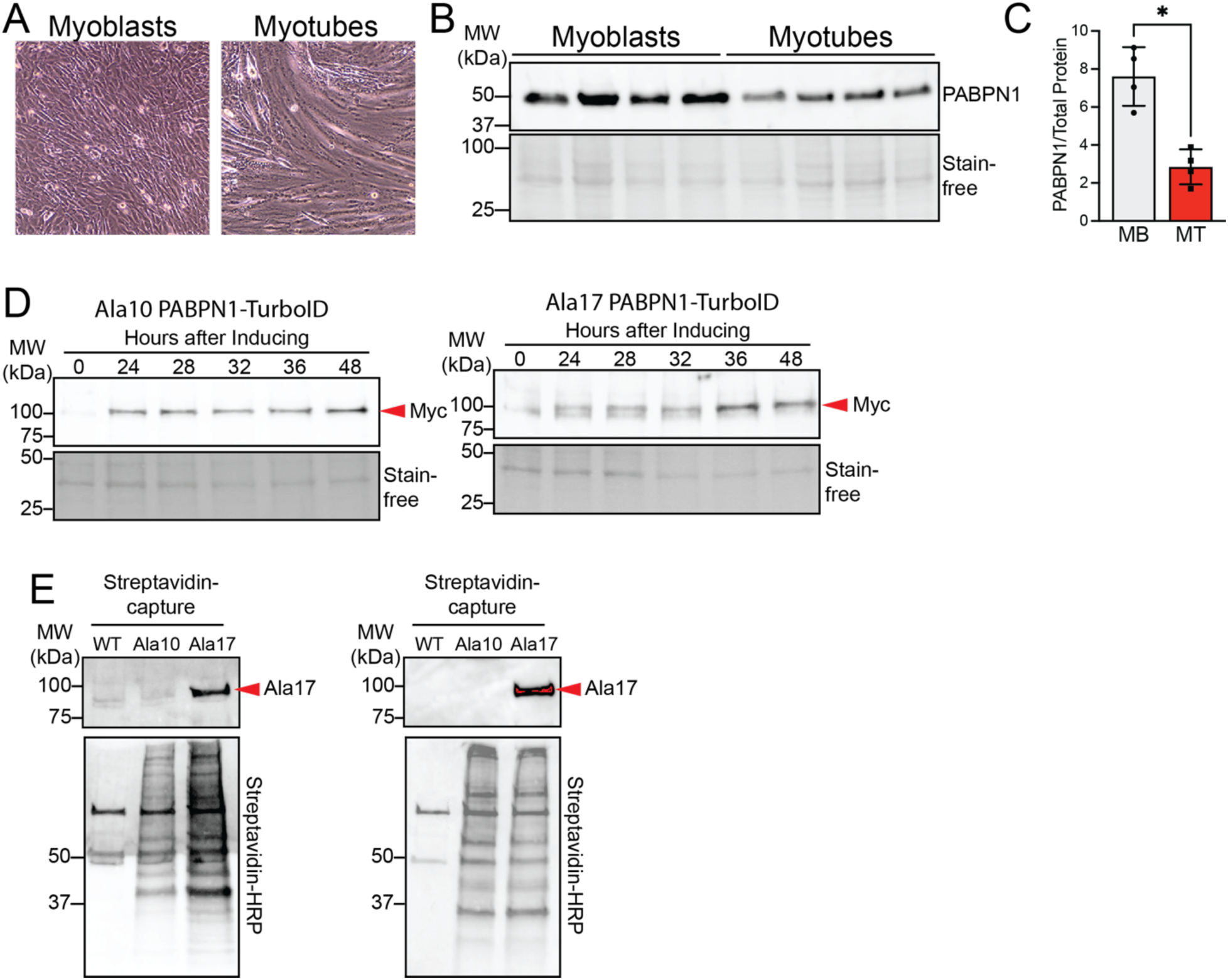
Additional characterization of PABPN1-TurboID constructs. **A)** Representative phase contrast image of C2C12 myoblasts versus myotubes. **B)** Immunoblot probed with an antibody to PABPN1 showing decreased PABPN1 in myotubes compared to myoblasts. **C)** Quantification of immunoblot in *B*. Shown is mean ± standard deviation for n = 4 experiments. Statistical significance determined using paired t-test. * p < 0.05. **D)** Immunoblots showing time course of doxycycline-induced expression of Ala10 (left) and Ala17 (right) PABPN1-TurboID constructs as detected using an antibody to the Myc tag. Stain-free imaging used as a loading control. **E)** Additional replicates of n2 (left) and n3 (right) blots of streptavidin elutions from Ala10 and Ala17 PABPN1-TurboID expressing myotubes probed with an antibody to the alanine expansion (top) to show alanine expansion in Ala17 PABPN-TurboID or probed with streptavidin-HRP to show increased presence of biotinylated proteins compared to wild type (WT) control myotubes.

**Supplemental Figure 2:**
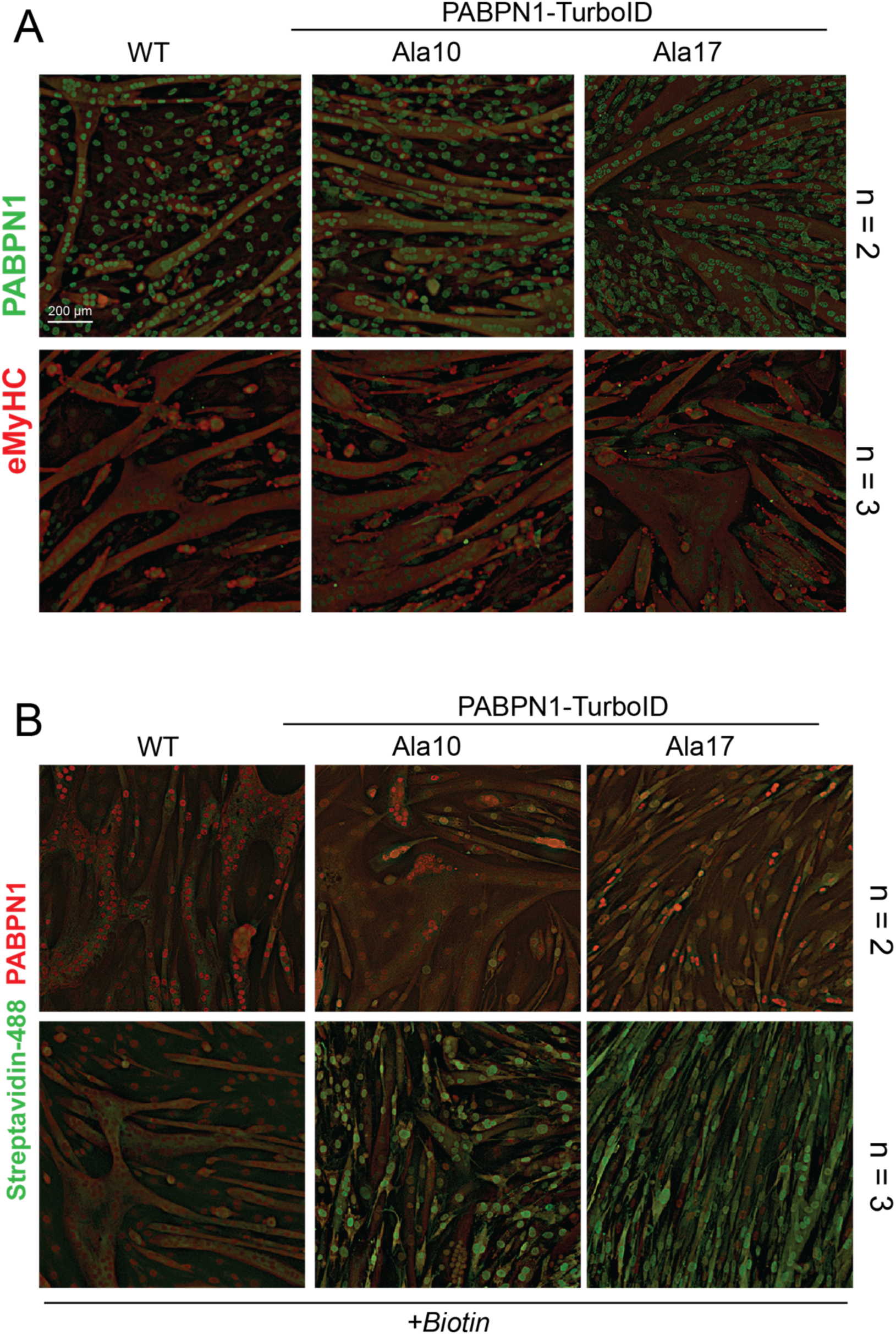
Additional replicates of myotubes stained for PABPN1 or for biotinylated proteins. **A)** Additional immunostaining of n2 (top) and n3 (bottom) with an antibody to PABPN1 (green) in myotubes from WT and stable cells expressing near-native levels of Ala10 and Ala17 PABPN1-TurboID. Immunostaining with an antibody to embryonic myosin heavy chain (red) was used to detect myotubes. **B)** Additional staining of n2 (top) and n3 (bottom) using AF488-conjugated streptavidin (green) to detect biotin and an antibody to PABPN1 (red).

**Supplemental Figure 3:**
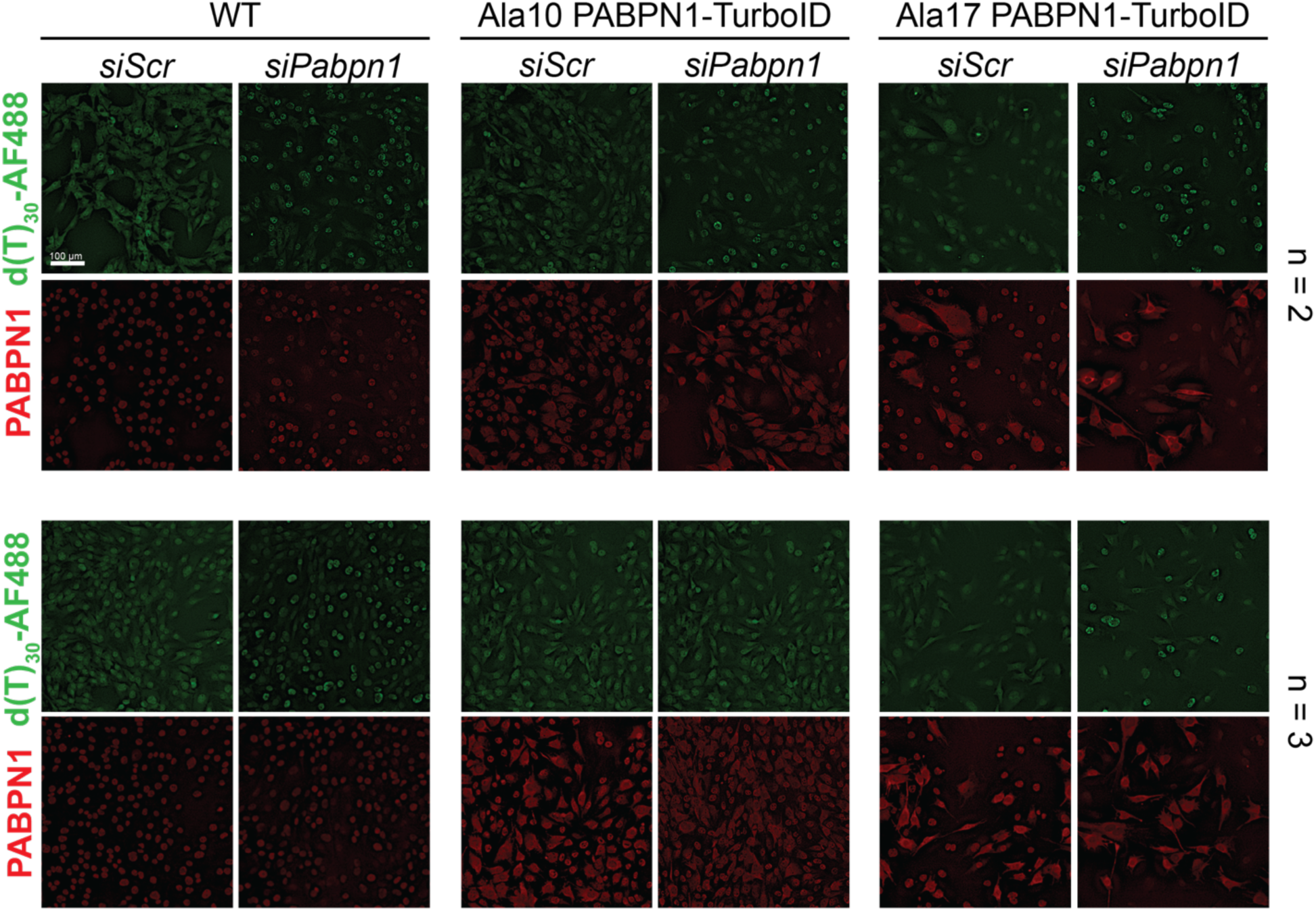
Additional replicates of d(T)_30_-AF488 FISH in *Pabpn1* knockdown myoblasts with and without expression of PABPN1-TurboID constructs. Additional d(T)_30_-AF488 FISH staining of n2 (top) and n3 (bottom) in *Pabpn1* knockdown in WT myoblasts or myoblasts expressing Ala10 or Ala17 PABPN1-TurboID.

**Supplemental Figure 4:**
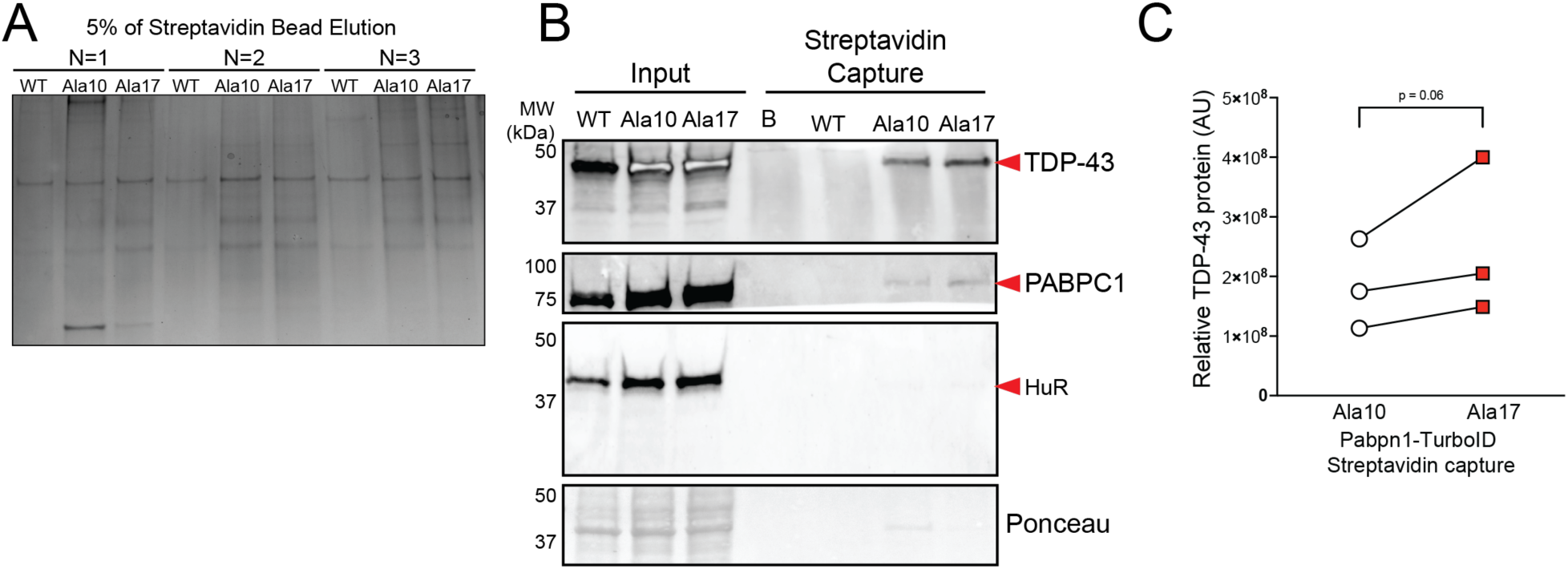
Preparation of PABPN1-TurboID expressing myotube lysates for proteomics. **A)** Silver-stained gel of 5% of streptavidin bead elutions from WT myotubes and myotubes expressing Ala10 or Ala17 PABPN1-TurboID. Shown are all three replicates used for comparative proteomics. **B)** Immunoblot of streptavidin elutions from WT myotubes and myotubes expressing Ala10 or Ala17 PABPN1-TurboID probed with antibodies to known PABPN1 binding partners TDP-43 and PABPC1 as well as HuR. Ponceau stain is used as a loading control. **C)** Quantification of TDP-43 band from streptavidin elution blot showing trending increase in TDP-43 detection in Ala17 PABPN1-TurboID proximal blot. Shown are n = 3 replicates analyzed by paired t test.

**Supplemental Figure 5:**
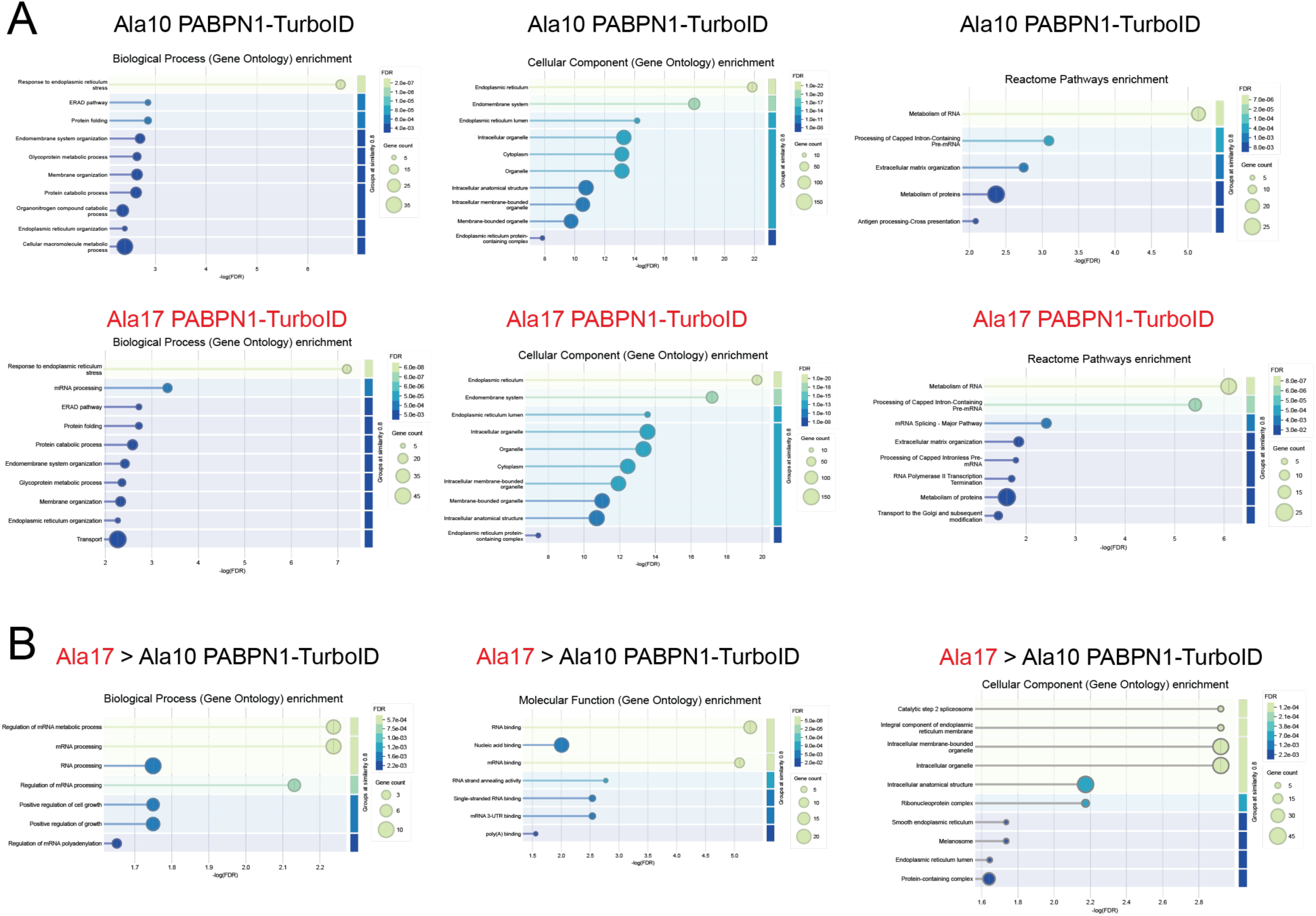
Additional functional annotation of proteomic data. **A)** Gene ontology biological process, cellular component, and Reactome pathways enriched in Ala10 PABPN1-TurboID (top) and Ala17 PABPN1-TurboID (bottom) proximal proteins. **B)** Gene ontology biological process, cellular component, and Reactome pathways enriched in Ala17 PABPN1-TurboID proximal proteins compared to Ala10 PABPN1-TurboID. In all cases, data shown are representative of proteins detected in at least two of three Ala10 or Ala17 PABPN1-TurboID replicates and not detected in WT controls.

**Supplemental Figure 6:**
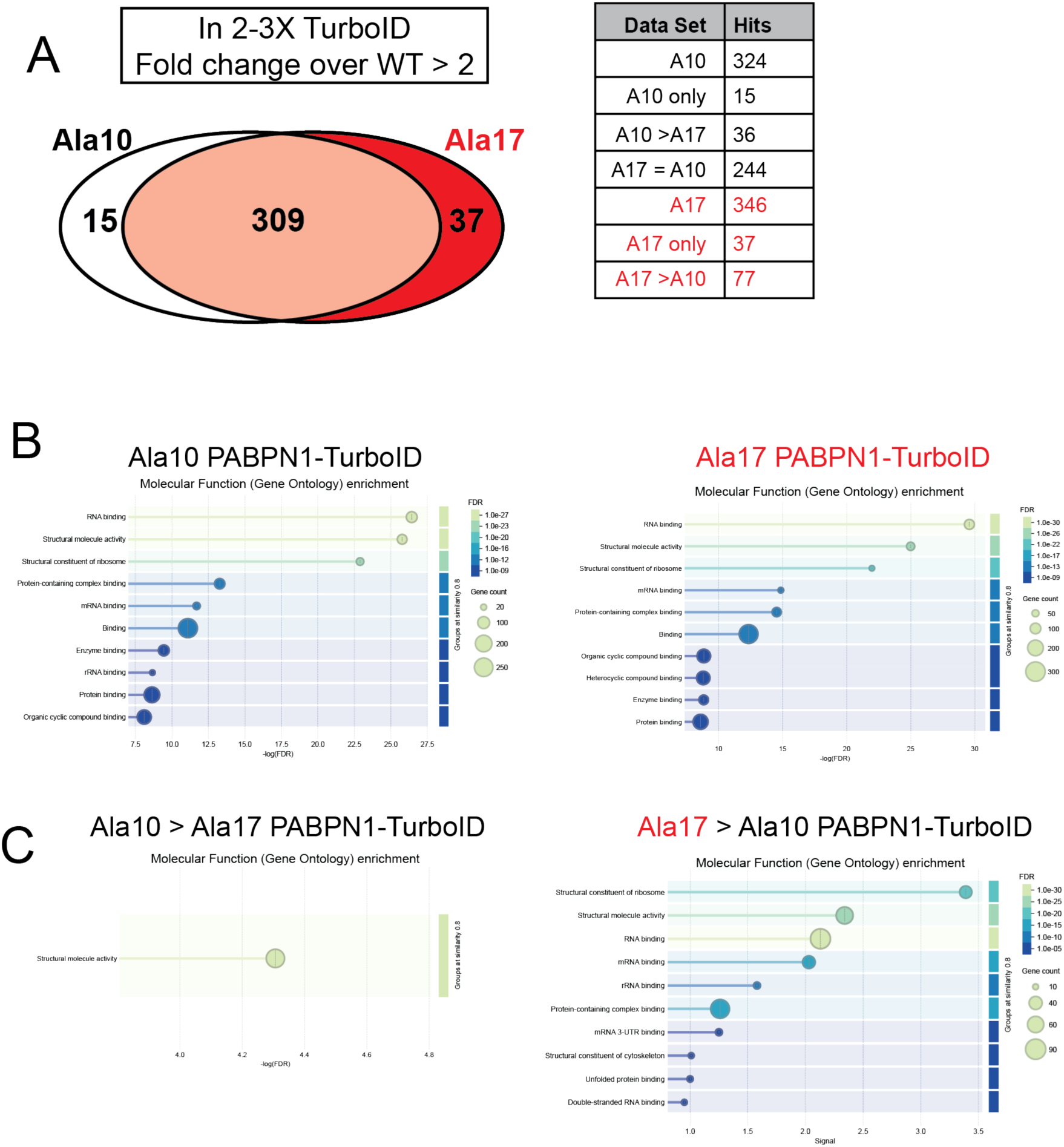
Functional annotation of more inclusive proteomic data. **A)** Proximal proteins detected in at least two of three PABPN1-TurboID replicates at least two-fold higher or more over WT controls. **B)** Gene ontology molecular function enrichment for Ala10 PABPN1-TurboID (left) and Ala17 PABPN1-TurboID (right) proximal proteins. **C)** Gene ontology molecular function enrichment for proteins enriched in Ala10 over Ala17 PABPN1-TurboID (left) and Ala17 over Ala10 PABPN1-TurboID (right).

**SUPPLEMENTAL TABLE 1.**
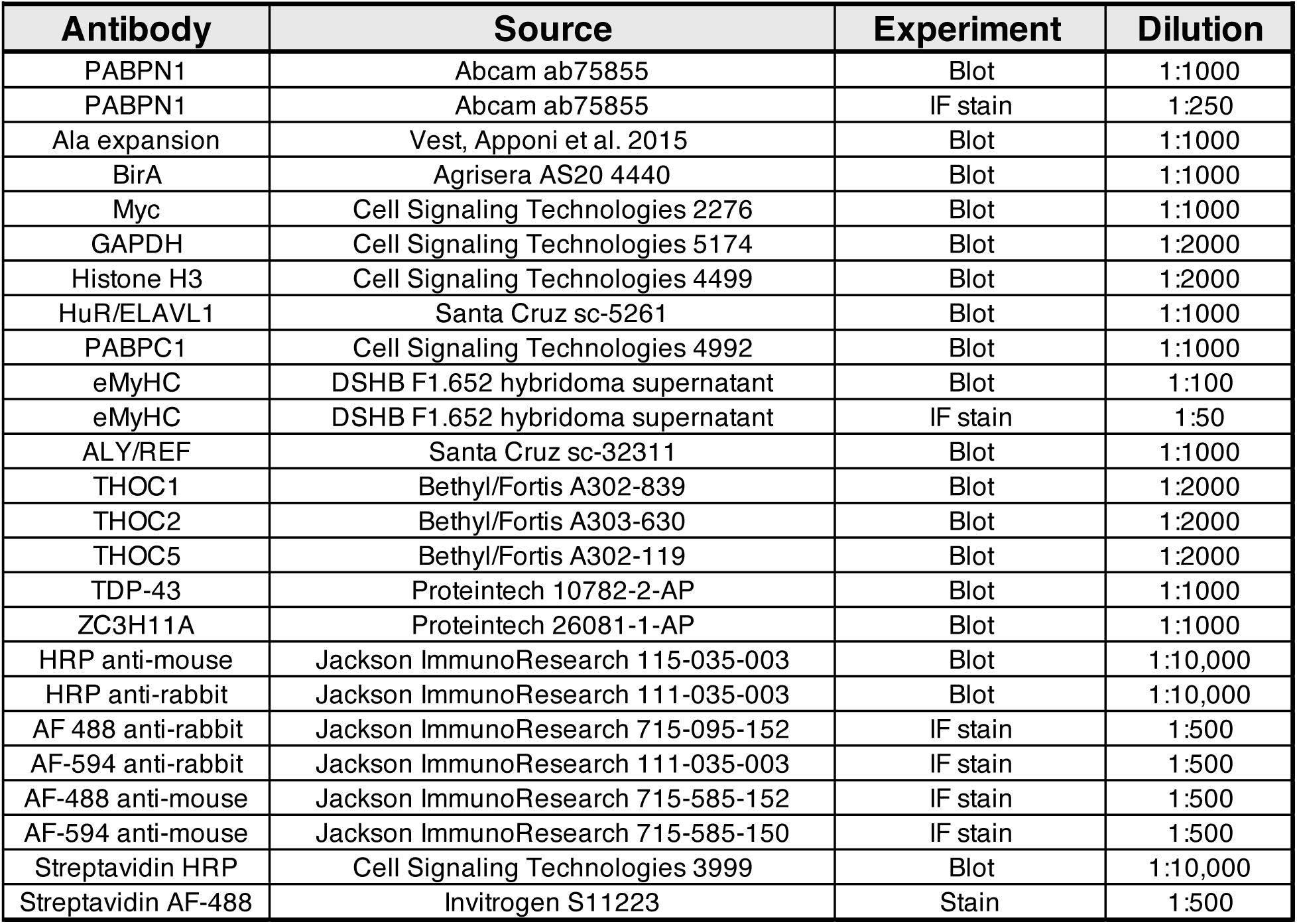

